# Surrogate selection oversamples expanded T cell clonotypes

**DOI:** 10.1101/2023.07.13.548950

**Authors:** Peng Yu, Yumin Lian, Cindy L. Zuleger, Richard J. Albertini, Mark R. Albertini, Michael A. Newton

## Abstract

Inference from immunological data on cells in the adaptive immune system may benefit from modeling specifications that describe variation in the sizes of various clonal sub-populations. We develop one such specification in order to quantify the effects of surrogate selection assays, which we confirm may lead to an enrichment for amplified, potentially disease-relevant T cell clones. Our specification couples within-clonotype birth-death processes with an exchangeable model across clonotypes. Beyond enrichment questions about the surrogate selection design, our framework enables a study of sampling properties of elementary sample diversity statistics; it also points to new statistics that may usefully measure the burden of somatic genomic alterations associated with clonal expansion. We examine statistical properties of immunological samples governed by the coupled model specification, and we illustrate calculations in surrogate selection studies of melanoma and in single-cell genomic studies of T cell repertoires.

**Funding:** This research was supported in part by the National Science Foundation (grant 2023239-DMS), and by grants from the National Institutes of Health: R01 GM102756, P01 CA022443, P01 CA250972, P50 CA278595, UL1 TR002373, P50 CA269011, and P30 CA014520. This work was also supported by resources at the William S. Middleton Memorial Veterans Hospital, Madison, WI, USA, and the UW Carbone Comprehensive Cancer Center. Additional support was provided by Ann’s Hope Foundation, Taking on Melanoma, the Tim Eagle Memorial, and the Jay Van Sloan Memorial from the Steve Leuthold Family Foundation, philanthropic support in the USA. The content is solely the responsibility of the authors and does not necessarily represent the official views of the NIH or the views of the Dept. of Veterans Affairs or the United States Government.

## 1. Introduction

### 1.1. Overview

With thymic-derived lymphocytes (i.e., T cells) sampled from peripheral blood or some other tissue compartment (e.g., tumor-infiltrating lymphocytes), any techniques that would enrich the sample for disease-relevant cells could be useful, considering the complexity of a typical T cell population and the potential for an improved understanding of the immune response to disease. For example, at writing we have no effective biomarkers to predict how a melanoma patient will respond to immune checkpoint inhibition therapy, though responses among similar patients may vary from morbid toxicity to full recovery (e.g., Ganesan and Mehnert, 2020; Shum, Larkin and Turajlic, 2022).

Surrogate selection restricts a lymphocyte sample *in vitro* to cells whose somatic ancestors had acquired and thus transmitted to them specific, selectable mutations. Selection assays based on mutations of the hypoxanthine-guanine phosphoribosyltransferase (HPRT) gene are most well studied, though the approach applies to any mutations that are neutral with respect to the immune response (Kaitz et al., 2022). As an immune-system probe, HPRT surrogate selection has been used to study a variety of environmental effects and disease processes (Albertini, Castle and Borcherding, 1982; Albertini, 2001; Kaitz et al., 2022). With continued focus on disease studies, we examine the sampling effects of surrogate selection; selected cells may represent *in vivo* amplified clones that are more likely to be disease relevant than clones of randomly sampled cells, and we seek a more thorough understanding of this enrichment phenomenon for the sake of improved experimental design and data analysis.

The idea that surrogate selection can enrich for clonally amplified T cells has provided a rationale in many studies, though quantitative treatments of this experimental-design strategy remain very limited. Statistical procedures have been deployed to test from sequence data the null hypothesis that enrichment is absent, and the mounting evidence supports the alternative (e.g., Pei et al., 2014; Zuleger et al., 2020). Considering cell growth dynamics, one would predict an increased prevalence of various somatic mutations in cells within an actively proliferating clone compared to a relatively quiescent one. Then conditioning on the presence of some such mutation in a sampled cell, Bayes’s rule would imply that the cell is more likely to be from the proliferating than the quiescent clone. Surrogate selection thus relies on the biological consequences of *in vivo* clonal proliferation to enrich for activated T cells in individuals with ongoing immunological response to disease. Understanding this enrichment effect is complicated by the enormous complexity of T cell population and properties of the distribution of clone sizes, but resolving these complications will inform investigations of surrogate selection as a mechanistic probe for fundamental biological/immunological processes. The main contribution of the present work is to quantify the enrichment effect of surrogate selection in an idealized but structurally relevant setting, and to leverage basic stochastic-process theory to confirm and characterize the enrichment phenomenon in this model. Our formulation also enables a study of distributional properties of elementary diversity statistics, of the type often used in experimental studies. We show that samples identified using surrogate selection have lower expected sample diversity, in agreement with empirical studies.

Our theoretical analysis exposes an interesting statistical prediction concerning somatic mutations that are unrelated to any selection assay. From contemporary single-cell genomic studies, we associate T cell clone sizes with estimates of somatic mutation burden, and thereby provide a new measure of somatic burden of a T cell receptor.

### 1.2. Immunological setting

Consider a person’s T cell repertoire, comprised of perhaps 10^11^ or more CD4+ and CD8+ naive, effector, and memory T cells, and partitioned into clonotypes within each of which the T cell receptor (TCR) sequence of the cells is constant (e.g., Nikolich-Žugich, Slifka and Messaoudi, 2004; Pennock et al., 2013; van den Broek, Borghans and van Wijk, 2018). The number of T cells in each clonotype fluctuates over time and usefully may be viewed as a stochastic process (Currie et al., 2012; Hodgkin, Dowling and Duffy, 2014; Desponds, Mora and Walczak, 2016; Gaimann et al., 2020; Smith et al., 2020; Molina-París and Lythe, 2021). Notably, a T cell receptor’s cognate antigen may induce cell division and expansion of the associated clonotype when appropriate costimulatory molecules are present. Complexity of the adaptive immune response warrants highly detailed stochastic-model dynamics, perhaps accounting for clonal competition or adaptation (e.g., Stirk, Molina-París and van den Berg, 2008; Lythe and Molina-París, 2018; Rane et al., 2018; Duque et al., 2020). However, even structurally simple models can support certain lines of investigation and can guide statistical analysis in the growing number of empirical studies. T cell receptor repertoire analysis has been critical in studies investigating antitumor responses as well as immune-related toxicity following treatment with immune-checkpoint blockade (e.g., Fairfax et al., 2020; Valpione et al., 2020; Lozano et al., 2022; Valpione et al., 2021).

### 1.3. Surrogate selection

In the absence of an assay to measure the proliferation history of a sampled T cell, surrogate selection provides an indirect measurement through the lens of neutral somatic mutation. The most well-studied case leverages an assay to score somatic mutations of hypoxanthine-guanine phosphoribosyltransferase (HPRT) (Albertini et al., 1990; Albertini, 2001). Other assays rely on an efficient approach to screen mutations in phosphoinositolglycan class A (PIG-A) genes (Peruzzi et al., 2010; Dobrovolsky et al., 2017). Coding an enzyme within the purine salvage pathway, HPRT normally helps to recycle nucleotide bases from degraded DNA. Its post-translational modifications also confer cytotoxicity to purine analogs, including 6-thioguanine (6TG). Cultured lymphocytes are thus unable to grow in the presence of 6TG unless they have incurred an inactivating HPRT mutation. Each surviving T cell in an HPRT assay reports that an HPRT mutation occurred in that T cell or in one of its somatic ancestors. The assay has been used to monitor somatic mutations in many settings, including, for example, in Chernobyl liquidators (Jones et al., 2002), in Iraq war veterans (Nicklas et al., 2015), and in studies of environmental exposures. Kaitz et al. (2022) reviews the implicit model for surrogate selection and the literature using HPRT surrogate selection in autoimmune diseases, cardiac transplantation, infectious diseases, a hematological disease, and cancer.

### 1.4. Summary of findings

The rationale for surrogate selection in disease studies is that it provides an enrichment for relevant T cell clonotypes. Some care is required in this argument, since while a large, expanded clonotype has higher sampling probability than any smaller clonotype, the vast diversity within a typical T-cell repertoire means that even large clonotypes remain a small fraction of the total population; indeed, most sampled cells come from small clonotypes. Basic stochastic process theory guides our effort to balance these factors. We find that if at any time point the vector of clonotype sizes in a repertoire is exchangeable, and if the temporal development of any one clonotype follows a sufficiently regular birth-death process, then surrogate selection via neutral somatic mutation enriches the sampled cells for those of larger clonotypes. We examine the impact of surrogate-selection on the expected value of sample diversity statistics. In empirical validations, we re-examine single-cell data from publicly available T cell repertoire samples that were obtained via 10x Genomics sequencing; in doing so we compute cell-level somatic burden statistics and associate this burden with clonotype size. We also review sample diversity statistics from available surrogate-selection studies.

## 2. One developing clonotype

### 2.1. Model set up

Our calculations begin by considering one clonotype of the many within an individual subject’s T cell repertoire. For definiteness, we label this clonotype *σ*, recognizing that *σ* resides in a large finite label set 𝒮, which we associate with the set of possible T cell receptor sequences. At time *t* ≥ 0 relative to some reference time point *t* = 0 (e.g., birth), clonotype *σ* consists of *N*_*σ*_(*t*) cells. If clonotype *σ* is ever non-empty, then there is some origin time, say *τ*_*σ*_, such that *N*_*σ*_(*t*) = 0 for *t < τ*_*σ*_ and *N*_*σ*_(*t*) *>* 0 only at times *t* ≥*τ*_*σ*_. We suppose that *N*_*σ*_(*τ*_*σ*_) = 1; that is, the clonotype originates upon successful completion of receptor-forming recombination events (Elhanati et al., 2018). After positive and negative selection induce thymocyte maturation, clonotype cells egress from the thymus and distribute themselves throughout the body; we expect this all occurs on a short time scale compared to the timing of typical observations, which might be from a mature subject’s peripheral blood or tumor-infiltrating lymphocytes, for example.

The stochastic process {*N*_*σ*_(*t*) : *t* ≥ 0} fluctuates in response to all sorts of cell-biological factors affecting cells in the clonotype, and must reflect a complex birth-death process (e.g., den Braber et al., 2012; Desponds, Mora and Walczak, 2016; Zhan et al., 2017). For example, in the presence of appropriate cytokines, T cell receptor interaction with cognate antigen triggers cell proliferation, while apoptotic signals can induce cell death. Our understanding of repertoire maintenance further supports the notion that if *N*_*σ*_(*s*) = 0 at time *s > τ*_*σ*_, then *N*_*σ*_(*t*) = 0 for all *t* ≥*s*. This is analogous to the infinite-alleles assumption in population genetics; here it means that a clonotype can only emerge once.

### 2.2. The branching tree

Following clonotype *σ* over time from *τ*_*σ*_, there is a series of event times at which cells in the clonotype either divide or die. Were we able to trace *σ*’s complete history, we would record a binary tree, such as in Figure 1. At some observation time *t*_obs_, each leaf of the tree is an extant cell that has experienced a number of cell divisions since *τ*_*σ*_. This division number is also called the depth of the leaf node. For a cell randomly sampled from the clonotype, let *D*_*σ*_ denote this division number; it has a probability distribution induced both by the stochastic development of *σ* and by the random selection of the extant cell. Fortunately, this distribution has been the subject of extensive study in the context of random binary trees (e.g., Lynch, 1965; Mahmoud, 1992; Aldous, 1996; Steel and McKenzie, 2001; Mahmoud and Neininger, 2003).

**FIG 1.**
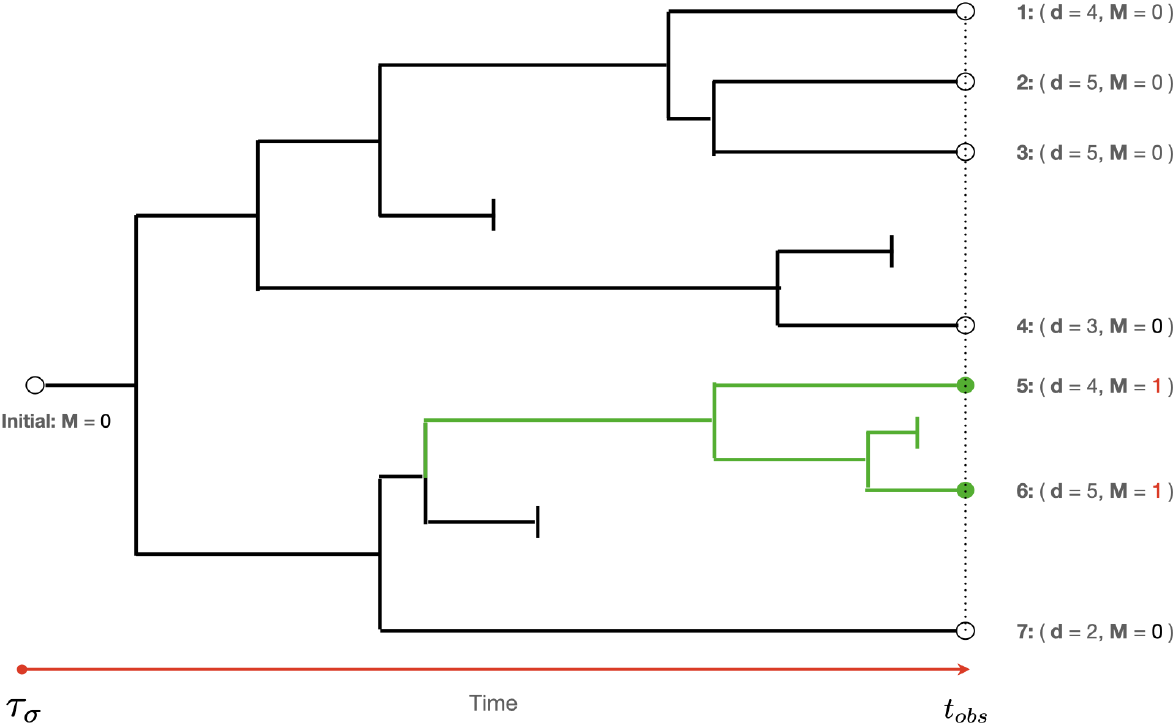
Binary tree formed by a developing clonotype, showing examples of cell division, cell death and mutation, and noting the number d of cell divisions experienced by each extant cell at time t_obs_. Green circles (extant cells 5 and 6) denote mutant T cells. Empty circles (1, 2, 3, 4 and 7) denote wild type T cells. Green lines denote evolution of mutant cells. Short vertical lines denote cell death.

In the Yule model for trees, each cell division acts on a random cell, as if by a pure-birth process without cell death. This symmetry over cell identity allows various explicit computations. In fact, the probability generating function (p.g.f.) of *D*_*σ*_ is

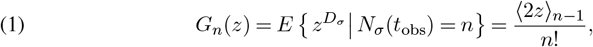

which is the formulation presented in (Mahmoud, 1992, Page 71-74), Eq. (2.4).^1^ Here ⟨*x*⟩ _*n*_ = *x*(*x* +1)(*x* +2) · · · (*x* + *n* − 1) is the rising factorial, which is conveniently expressed in terms of Gamma and Beta functions Γ and *B* as:

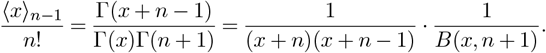

The p.g.f. *G*_*n*_ helps us connect the T cell repertoire with surrogate-selection dynamics. Before pursuing that calculation, we note that the expectation and variance of *D*_*σ*_ are also available, with both well approximated by twice the natural logarithm of *n*, and that as *n* increases, 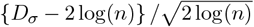 converges in distribution to a standard normal variate (Brown and Shubert, 1984; Mahmoud and Neininger, 2003). Roughly, a randomly sampled cell from a randomly proliferating clonotype of current size *n* (and ignoring cell death) has experienced about 2 log(*n*) cell divisions since receptor formation in the thymus. Sampling from the conditional distribution of *D*_*σ*_ |*N*_*σ*_(*t*_obs_) = *n* is reported in Figure 2, revealing this proliferation effect for a handful of clonotype sizes. For completeness, we note the p.m.f. of *D*_*σ*_ is, as derived in Lynch (1965),

**FIG 2.**
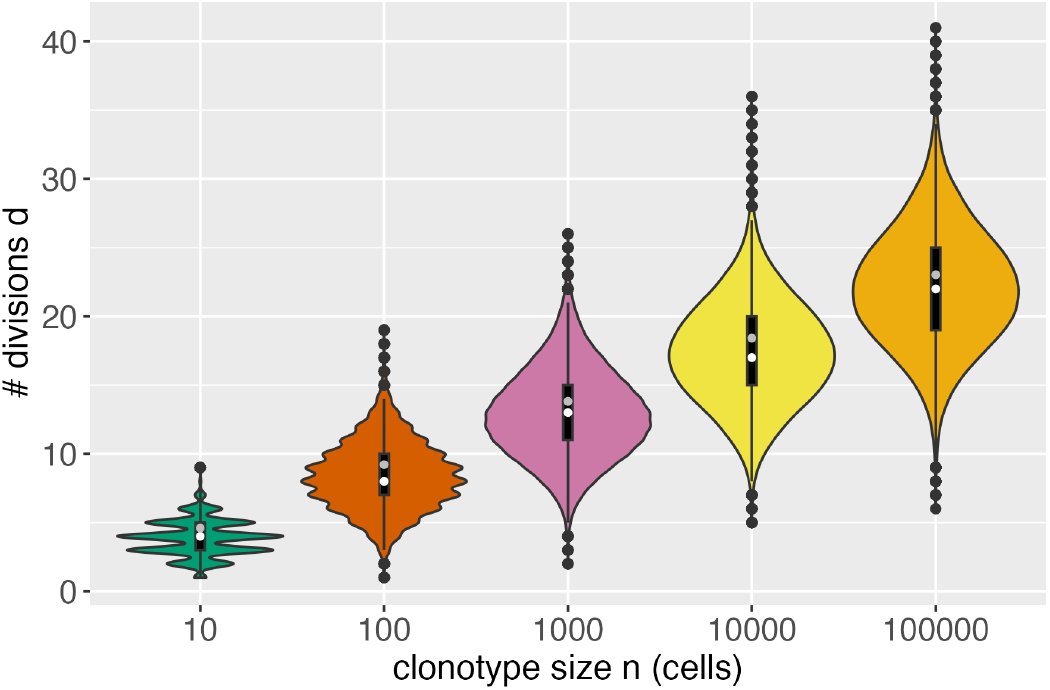
Proliferation effect: Shown are violin plots of the division number D_σ_for cells in randomly developed binary trees, having various sizes, n, at observation time. We used **R** packages **ape**, to simulate Yule trees, and **adephylo**, to count divisions (Paradis and Schliep, 2019; Jombart, Balloux and Dray, 2010). Each plot summa-rizes 100,000 simulated D_σ_values. Empirical medians (white) and asymptotic means 2 log(n) (grey) are shown.

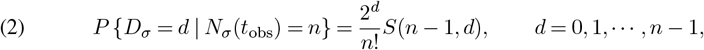

where *S*(*n* − 1, *d*) is the unsigned Stirling number of first kind.

### 2.3. Neutral mutations

Surrogate selection aims to use neutral genomic mutations – mutations that do not affect clonotype growth dynamics – as probes to report on these very same dynamics. Uncorrected mitotic errors or other mutagenic effects are expected to occur at some rate throughout the developing repertoire. We focus on mitotic mutations that affect a single daughter cell, that are irreversible, and that occur independently across cell divisions. Less prevalent mechanisms may induce mutations in both daughter cells (e.g., double-stranded breaks) or separately from mitosis (e.g., ionizing radiation), and statistical formulations may be adapted to these cases (e.g., Kendall, 1960; Roshan, Jones and Greenman, 2014). We use *θ* ∈ (0, 1*/*2) to denote the relative frequency of mutations at a given locus (e.g., HPRT) per daughter cell; i.e., 2*θ* is the mutation frequency per cell division.

Consider the thought experiment to sample a single cell uniformly at random from the extant clonotype *σ* at time *t*_obs_, and let *M*_*σ*_ be the binary (0*/*1) indicator that the sampled cell harbors a mutation at the locus in question. We recognize that *M*_*σ*_ really indicates that a mutation event occurred somewhere in the ancestral lineage of the cell, and thus

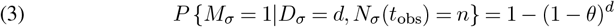

where *D*_*σ*_ is the division number for this random cell. (The cell is not mutant if none of the *d* opportunities for mutation yield such.) Incidentally, (3) implies that *M*_*σ*_ and *N*_*σ*_(*t*_obs_) are conditionally independent given *D*_*σ*_. Our first finding concerns the rate of mutant genotype in clonotypes of a given size, and is obtained by marginalizing the distribution of *D*_*σ*_. With neutral mutations in a Yule tree model, define *ψ*_*n*_ := *P* {*M*_*σ*_ = 1|*N*_*σ*_(*t*_obs_) = *n*}, and note,

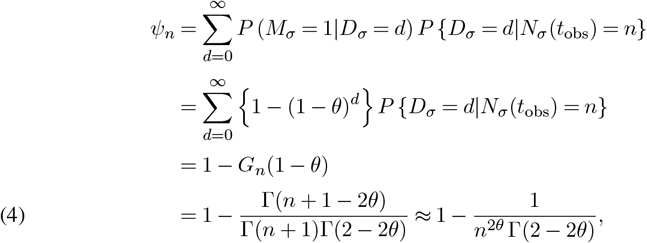

with the approximation on the last line improving for increasing *n*. Result (4) quantifies the intuition that proliferating clonotypes provide a greater number of chances for mutation. With *θ >* 0, lim_*n*→ ∞_ *ψ*_*n*_ = 1, and so an ever-proliferating clonotype is eventually dominated by mutant cells. This matches limit theory for birth-death processes in which the growth rate of mutant cells is no less than that of wild-type cells (e.g., Cheek and Antal, 2018).

We are not too concerned with the total number of mutant cells in the clonotype, whose expected value is *n* time the per cell rate in (4), though our diversity calculations in Section 3.5 rely on this distribution. That total mutant count is interesting in other settings, and is governed by the Luria-Delbrück distribution; see Angerer (2001) or Roshan, Jones and Greenman (2014) for the exact, non-asymptotic formulation. The reader may check that our formula (4) matches the first-moment formula from Roshan, Jones and Greenman (2014), Theorem 3.3, taking *n* = *k* and *μ*_1_ = 1 − *μ*_0_ = 2*θ*; interestingly, a quite different approach is taken in that paper.

### 2.4. Enrichment and Bayes rule

The development so far has emphasized probabilities that condition in some way on clonotype size. Next we layer in a distribution on that size itself; the stochastic evolution of a specific clonotype *σ* induces a distribution on the size *N*_*σ*_(*t*_obs_) at observation time. For example, the linear pure-birth model leads to the Geometric{exp(−*λ*_*σ*_*t*_obs_)} distribution,

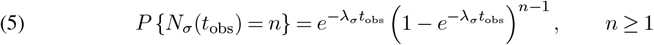

where *λ*_*σ*_ is the birth rate (rate of cell division). Further, compounding over *λ*_*σ*_ gives the Yule-Simon law, with parameter *ρ >* 0,

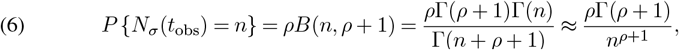

where the approximation improves with increasing *n*. This is approximately a power-law, or Zipf distribution, which has been found to fit many T-cell repertoires (e.g., Bolkhovskaya, Zorin and Ivanchenko, 2014; Desponds, Mora and Walczak, 2016; Koch et al., 2018; Gaimann et al., 2020; de Greef et al., 2020), with exponents *ρ* in the range 0.05 to 0.2. Other marginal distributions on *N*_*σ*_(*t*_obs_) may be induced by more complex stochastic dynamics, such those modeling competition and thymic pressure (Lythe and Molina-París, 2018).

Combining the forward, mutant-genotype model (4) with a size model *P* {*N*_*σ*_(*t*_obs_) = *n*}, we have by conditioning:

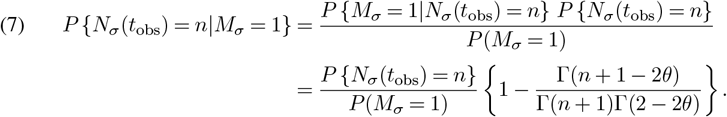

This Bayesian inversion of (4) quantifies surrogate selection’s enrichment effect in the pure-birth case. One setting is shown in Figure 3, which illustrates the suppression of probability on small clonotypes and inflation for larger ones. In that example, the median of the unconditional Geometric distribution is 6931 cells, while after conditioning on *M*_*σ*_ = 1, the median clonotype size shifts up to 8139 cells. This effect is not limited to the marginal Geometric law. Figures 4 show the result for a Logarithmic distribution (p.m.f. proportional to *p*^*n*^*/n*) and a Yule-Simon law (6), respectively. Summarizing the findings for a single, developing clonotype, we have:

**FIG 3.**
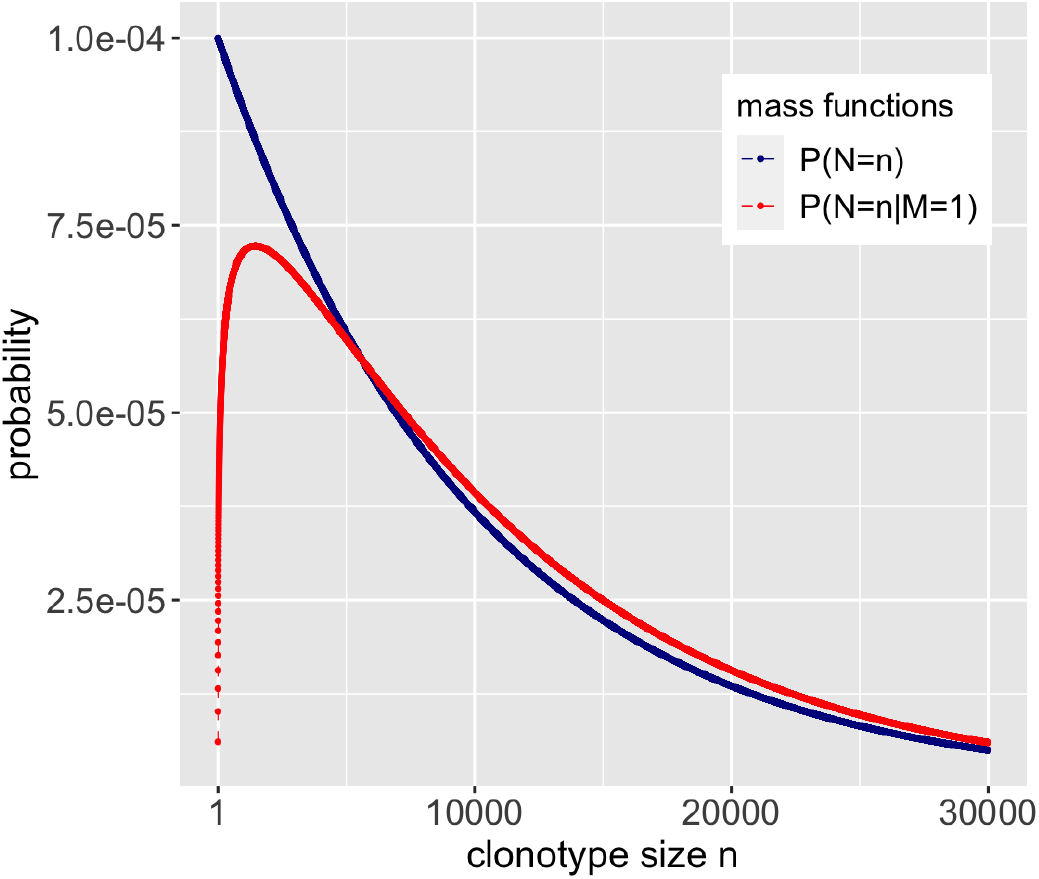
*P* {*N*_σ_ (*t*_obs_) = *n*|*M*_*σ*_ = 1} (red) when the marginal distribution (blue) is a Geometric distribution with parameter 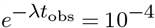 and the mutation frequency θ = 10^−6^. The crossover point n_cross_is 5624 cells.

**FIG 4.**
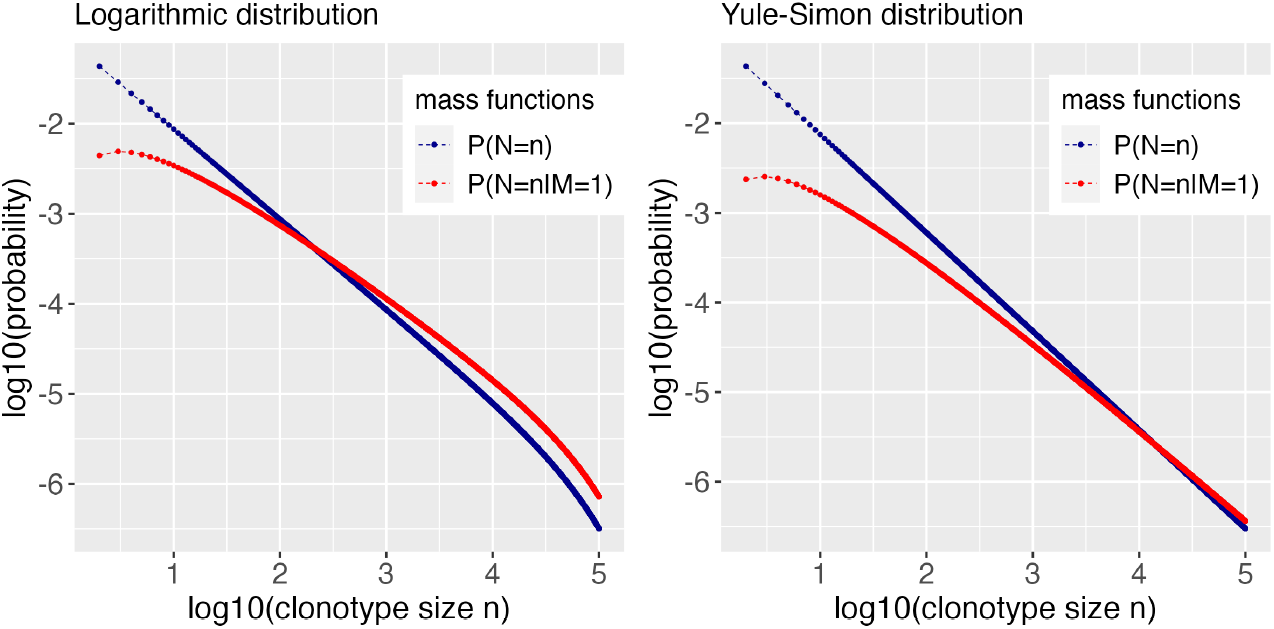
*P* {*N*_*σ*_ (*t*_obs_) = *n*|*M*_*σ*_= 1} (red) when the marginal clonotype size distribution (blue) is a Logarithmic distribution (left) or a Yule-Simon distribution (right), with parameters p = 1 − 10_−5_for Logarithmic distribution and ρ = 0.1 for Yule-Simon distribution. Mutation frequency θ = 10^−6^in both cases. The crossover point n_cross_ equals to 326 cells under Logarithmic distribution, and n_cross_= 14270 under Yule-Simon distribution.

proposition 1. *Suppose that, regardless of the marginal distribution of N*_*σ*_(*t*_obs_), *each cell division in the developing clonotype σ increases the clonotype size by 1 and occurs on a random extant cell, that a non-mutant dividing cell produces one mutant descendant (w.p*. − 2*θ) or no mutant descendants (w.p*. 1 − 2*θ), that descendants of a mutant dividing cell are both mutants, that there are no cell deaths, and that σ began with a single non-mutant cell. If M*_*σ*_ *indicates that a randomly sampled cell from σ at time t*_obs_ *is mutant, then the enrichment ratio f*_*n*_ := *P* {*N*_*σ*_(*t*_obs_) = *n*|*M*_*σ*_ = 1} */P* {*N*_*σ*_(*t*_obs_) = *n*} *is:*

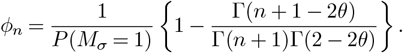

*Further, f*_*n*_ *is strictly increasing and approaches* 1*/P* (*M*_*σ*_ = 1) *>* 1 *as n* → ∞.

Two immediate corollaries assure that: (1) there exists a crossover point *n*_cross_ with *ϕ* _*n*_ *<* 1 when *n < n*_cross_ and *ϕ* _*n*_ *>* 1 when *n > n*_cross_, and (2) the conditional distribution is stochastically larger than the marginal distribution, which is another perspective on the notion that mass is pushed towards larger clonotypes. In fact, monotonicity of *ϕ* _*n*_ amounts to saying that the marginal and conditional distributions satisfy the monotone likelihood ratio ordering, which is stronger than stochastic ordering of c.d.f.’s: {*P N*_*σ*_(*t*_obs_) ≥*n*| *M*_*σ*_ = 1} ≥*P* {*N*_*σ*_(*t*_obs_) ≥*n*} (see Pfanzagl, 1964). Among other things, it also follows that the conditional distribution of *N*_*σ*_(*t*_obs_) given *M*_*σ*_ = 1 has larger expected value than the marginal distribution. Conceptually, learning that the sampled cell is mutant tells us that the clonotype is probably larger than we would have guessed otherwise.

### 2.5. Beyond pure birth

Relaxing the no-cell-death assumption makes quantifying enrichment more difficult. Explicit calculations in one example (Appendix A) show that conditioning on *M*_*σ*_ = 1 does not necessarily enrich for larger clonotypes. That highly stylized example captures features of clonal expansion followed by rapid clonal decline. The intuition is that having sampled a mutant cell, we may only know that its containing clonotype is relatively old, rather than knowing this clonotype is relatively large. These two features are equivalent in the pure-birth model. To develop this intuition further, we pursue calculations in a well-behaved but general class of birth-death processes, and we find conditions within this class which assure the enrichment-for-larger-clonotypes phenomena.

At times *τ* _1_ *< τ* _2_ *<* · · ·after *τ*_*σ*_, changes *A*_1_, *A*_2_, · · · occur that either increase the clonotype size (*A*_*i*_ = 1) or decrease the clonotype size (*A*_*i*_ = −1), in the first case by division of a random cell, and in the latter by death of a random cell. Then at time *t*, the clonotype size 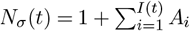 where τ _*I*(*t*)_ ≤ *t < τ* _*I*(*t*)+1_. We suppose this size process *N*_*σ*_(*t*) is not explosive, and thus only a finite number of *τ*_*j*_’s can occur in any finite time interval. We ask that *A*_*i*_ be independent of event times *τ*_1_ *< τ*_2_ *<* so that the discrete clonal history may be treated separately from questions of temporal rates of change. Further, we do not require a Markov condition, though we are mindful that having *A*_*i*_ conditionally independent of past changes given 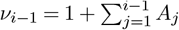 provides for a Markovian jump chain *ν*_*1*_, *ν*_*2*_, · · ·, with *N*_*σ*_ (*t*) = ν _*I*(*t*)_ (e.g., Grimmett and Stirzaker, 2001, pg 265). Considering mutation status along the jump chain, we introduce

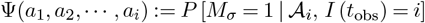

where 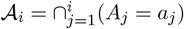 tracks the specific birth-death steps; thus Ψ is the conditional mutant frequency of a cell sampled from *σ* just after the *i* birth-death steps indicated by 𝒜_*i*_. Obviously we cannot sample a cell from an empty clonotype, so we furthermore condition on non-extinction, i.e. *ν*_*i*_ ≥ 1 for all *i*. The Ψ function generalizes the pure-birth *ψ*_*n*_ sequence (4), which we recover with *i* = (*n* − 1) and all *a*_*j*_ = 1, for example.

proposition 2. *In a birth-death process as defined above, Z*_*i*_ := Ψ(*A*_1_, *A*_2_, · · ·, *A*_*i*_) *is non-decreasing in i. If with probability one* 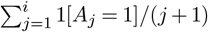 *diverges as i* → ∞, *then Z*_*i*_ *converges almost surely to the limit* 1, *and also E*(*Z*_*i*_) = *P* [*M*_*σ*_ = 1|*I*(*t*_obs_) = *i*] *converges to 1. Additionally, if ξ*_*n*,*i*_ := *E*(*Z*_*i*_|*ν*_*i*_ = *n*) *is non-decreasing in i* ∈ {*n* − 1, *n* + 1, *n* + 3, · · ·} *for each n, then P* [*M*_*σ*_ = 1|*N*_*σ*_(*t*_obs_) = *n*] ≥ *ψ*_*n*_.

In a linear birth-death process for example, and ignoring extinction for the moment, the *A*_*i*_’s are i.i.d., with *P* (*A*_*i*_ = 1) = *λ/*(*λ* + *μ*) for birth rate *λ >* 0 and death rate *μ* ≥ 0. It is well known that extinction is almost sure when *λ* ≤ *μ*, but also that extinction occurs with probability *μ/λ* as long as *λ > μ* (e.g., Grimmett and Stirzaker, 2001, pg 272). We would meet the requirements of Proposition 2 in this case; conditioning on non-extinction conditions on an event of positive probability. Note too that the divergence requirement follows immediately from the three-series theorem (e.g., Billingsley, 1995, pg 290). We have a recursive formula for *ξ*_*n*,*i*_ = *E*(*Z*_*i*_|*ν*_*i*_ = *n*); namely under the Markov condition for *ν*_1_, *ν*_2_, · · ·,

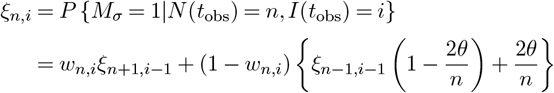

where *w*_*n*,*i*_ = *P* (*A*_*i*_ = −1| *ν*_*i*_ = *n*). We have not identified conditions assuring this *ξ*_*n*,*i*_ sequence is non-decreasing in *i* for each *n* (a requirement for Proposition 2); but numerical experiments in the linear birth-death model (Figure S1) give us confidence that this condition holds in relevant settings. The final lower-bound result in Proposition 2 means that conditioning on mutant status does enrich for larger clonotypes, thus extending Proposition 1. In any case, the monotonicity of *E*(*Z*_*i*_) indicates that such conditioning enriches for older clonotypes regardless of properties of *ξ*_*n*,*i*_.

## 3. Sampling from the repertoire

### 3.1. Model set up and size bias

Calculations so far refer to the random development of a single clonotype and its internal mutation rate. More relevant to experimental data are calculations that allow for sampling from the full repertoire, and thus the simultaneous development of many clonotypes. We eschew detailed, cell-biological considerations, though we do provide necessary structural elements to allow for a distributional comparison of diversity statistics computed either from wild type or mutant T cell fractions. First we address a curious size-biased sampling effect that emerges in considering the full repertoire, in contrast to the single clonotype from Sections 2.4 and 2.5.

We focus on a single observation time *t*_obs_, at which point the repertoire 𝒮 is comprised of non-empty clonotypes 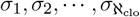, of sizes 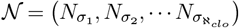, with 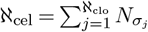 equal to the overall number of cells in the repertoire. We treat ℵ_clo_ and ℵ_cel_ as large constants, and, considering this snapshot of the repertoire, here we appreciate but do not emphasize with notation anything about the temporal, stochastic development of the clonotypes; for instance we ignore the multitude of receptors that are not extant at *t*_obs_, and we therefore have 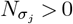 for all *j*. We allow that some more primitive generative stochastic process may underlie the clonotype counts, but we focus on their conditional joint distribution given the total number of cells ℵ_cel_ and the total number of extant clonotypes ℵ_clo_, which in adult humans may be on the order of 10^11^ and 10^8^, respectively. The same technical device was used by Rothman and Templeton (1980) in studying statistical properties of other assemblages, where additionally the assumption of finite exchangeability is helpful in revealing interesting system properties. We also adopt the finite exchangeability assumption for the joint mass function,

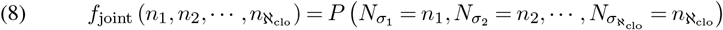

for counts *n*_*j*_ ≥ 1, which not only simplifies the specification, but also means that joint probability masses depend on the frequency spectrum holding the *counts-of-counts*: *C*(*k*) = ∑_*σ*_ 1[*N*_*σ*_ = *k*]. Figure 5 realizes a small synthetic example.

**FIG 5.**
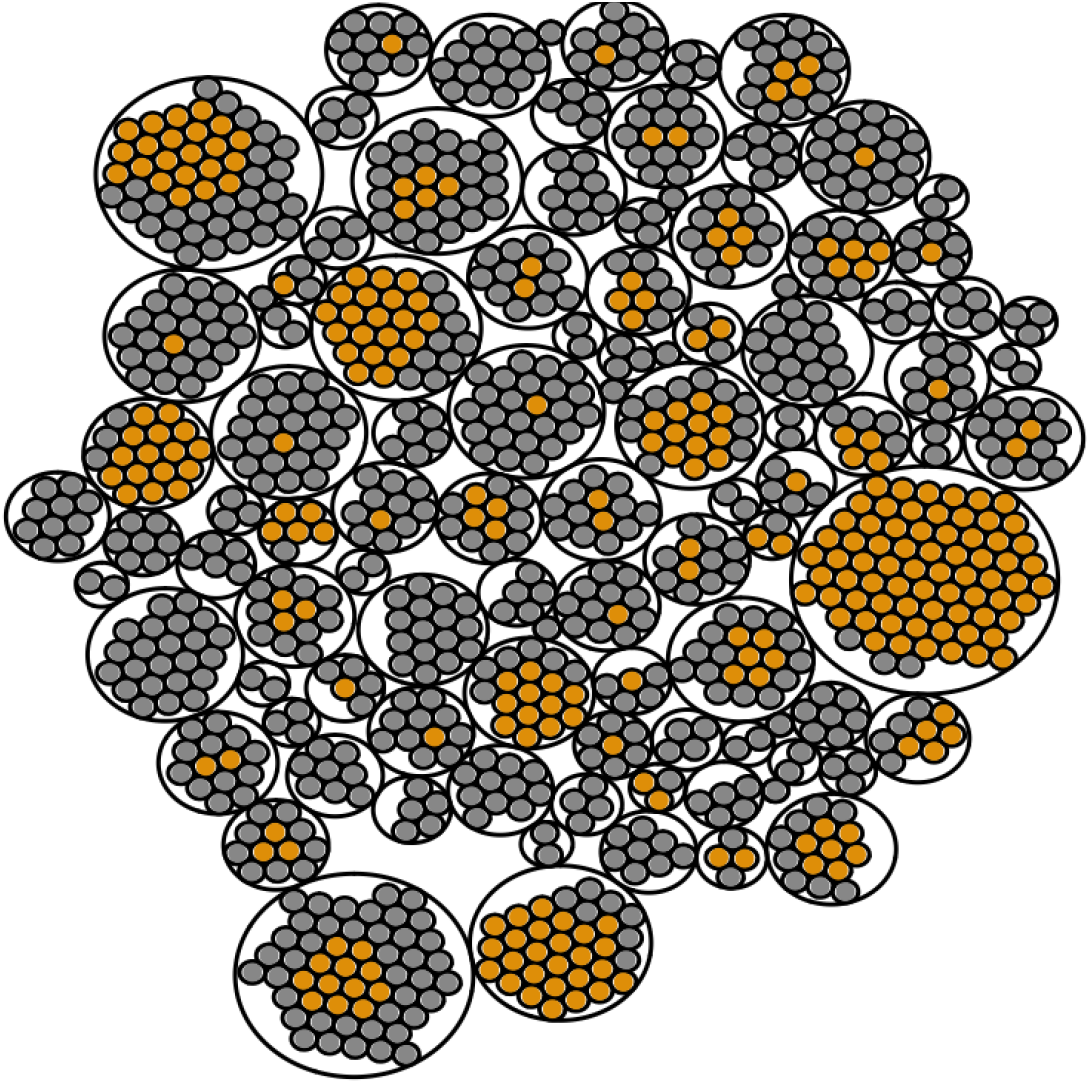
Simulated repertoire of ℵ_cel_ = 1000 cells comprising ℵ_clo_ = 100 non-empty clonotypes (encasing circles). The 287 mutant cells are orange/rust, and the remaining 713 wild-type cells are grey, giving a realized mutant frequency 0.287. As predicted mathematically, the larger clonotypes have an over-representation of mutant cells. Sampling uniformly among clonotypes, the average extant clonotype size is 10.0 cells; given the sampled clonotype contains a mutant cell, the average clonotype size is 16.0 cells. On the other hand, sampling uniformly among cells, the average clonotype size of the sampled cell (i.e., with size bias) is 23.0 cells. The average clonotype size when sampling mutant cells, however, is even larger, at 27.7 cells. This synthetic data was simulated from a Bose-Einstein clone-size model and a Luria-Delbrück mutation model, with mutation frequency θ = 0.05.

To appreciate the size-bias issue, consider sampling a single cell uniformly from the reper-toire, and let *S* ∈ 𝒮 denote its clonotype identifier. We recognize that *N*_*S*_, the size of the clonotype holding the sampled cell, is random owing to both the random development of the repertoire, as governed at least at the observation time by (8), and owing to the sampling of a cell from the repertoire. Under exchangeability, for *n* ≥ 1:

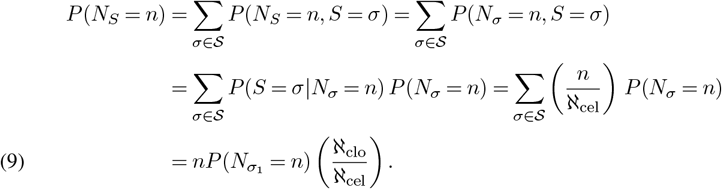

Size bias is reflected in the multiplication by *n* in (9). It conveys the fact that sampling a cell uniformly at random from a randomly developing repertoire is different (i.e., is biased towards larger clonotypes) than sampling a cell uniformly at random from a randomly developing clonotype. In any case, surrogate selection aims to further bias distributions towards larger clonotypes than would be obtained marginally. Before studying this enrichment, it is helpful to investigate a few exchangeable models and their relationship to well-known marginal distributions.

### 3.2. Joint assemblages and limiting margins: examples

By various compounding and conditioning operations applied to a collection of independent Poisson variates, Rothman and Templeton (1980) obtained an interesting exchangeable specification that we reconsider for (8):

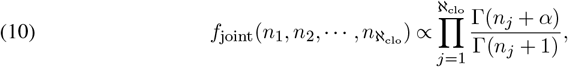

where the system-defining parameter *α >* 0 reflects dynamics of the assemblage. By modifying limiting regimes for ℵ_cel_, ℵ_clo_, and *α*, Rothman and Templeton (1980), *inter alia*, recovered reference marginal distributions distinguished especially by tail behavior. For example, setting *α* = 1 is the Bose-Einstein case. Sending ℵ_clo_*/*ℵ_cel_ *γ*_0_ (0, 1) as both the numerator and denominator diverge in this case, the marginal limiting distribution of any one clonotype size is Geometric(*γ*_0_), as in (5), which matches the pure-birth Yule tree model, with 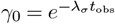. Similarly, if *α* → 0, the limiting margin is the Logarithmic distribution, with p.m.f. proportional to 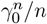; and if the limit of ℵ_clo_*/*ℵ_cel_ itself has a Beta(*ρ*, 1) distribution, then the limiting margin is the Yule-Simon power law (6). Empirical size distributions from the Bose-Einstein simulation conform nicely to these theoretical predictions (Figure S3). These intriguing relationships provide a modeling framework allowing us to elaborate single-clonotype calculations (Section 2) into the context of full-repertoire sampling. In particular, where various conditions on the joint assemblage give rise to different limiting marginal distributions for a given clonotype’s *N*_*σ*_, we can similarly deduce the size-biased distribution of *N*_*S*_. Details are provided in Appendix B; summarizing here, the size-biased version of the Geometric (5) has p.m.f. 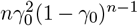, and the size-biased version of the Yule-Simon (6) has the p.m.f. *ρnB*(*n, ρ* + 2); see also Fig S2. We are not using these distributions for any sort of model-based inference from data; rather, we are exercising them primarily to explore implications of single versus multi-clonal analysis.

### 3.3 Enrichment

Size bias attributable to repertoire versus single-clonotype sampling does not alter the basic enrichment properties revealed in Propositions 1 and 2, except for a slight change in constants. For example, with the mutation model as in Section 2.4, and such that within each clonotype the stochastic process meets the conditions of Proposition 1, we have:

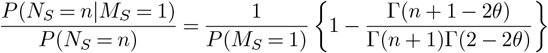

which is also a strictly increasing function of *n* that approaches limit 1*/P* (*M*_*S*_ = 1). The result follows from the single-clonotype sampling result (4), Bayes’s rule, and the equality:

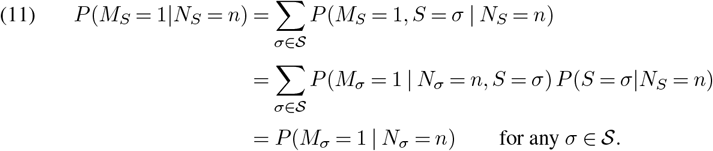

By analogy, Proposition 2 may also be extended to sampling from the full repertoire. In summary,

proposition 3. *If clonotype sizes at observation time t*_obs_ *are exchangeable, as in (8)*, *and if each individual clonotype evolves to its size at t*_obs_ *according to the dynamics in Proposition 1 or Proposition 2*, *then conditional on mutation M*_*S*_ = 1 *of a cell randomly drawn from the full repertoire, the enrichment ratio P* (*N*_*S*_ = *n M*_*S*_ = 1)*/P* (*N*_*S*_ = *n*) *eventually exceeds 1 for sufficiently large n*.

The enrichment phenomenon is illustrated in the synthetic repertoire in Figure 5, which shows mutant and wild-type subclones of various clonotypes, and highlights how sampling the mutant fraction would bias towards larger clonotypes.

### 3.4. Mutant Frequency

A random cell from the repertoire is more likely to be mutant than a random cell from any specific, randomly developing clonotype: *P* (*M*_*S*_ = 1) *> P* (*M*_*σ*_ = 1), which we confirm in the Appendix C by a calculation similar to (9). This mutant frequency *P* (*M*_*S*_ = 1) is of independent interest, and can be estimated by various dilution assays. As reviewed in Kaitz et al. (2022), the mutant frequency is different from the mutation frequency *θ*. The former considers the rate at which mutant cells are found in a sample from the repertoire; the latter is the rate that mutations emerge among cell divisions in a developing clonotype. Table S2 offers some numerical results for the Bose-Einstein assemblage.

### 3.5. Diversity statistics

An important motivation for the preceding theoretical calculations is to understand the impact of surrogate selection on statistics from a random sample from a repertoire. Suppose the amount of sampled material from one subject is a fraction ϵ = *n*_samp_*/*ℵ_cel_ of the entire repertoire, and let *X*_*σ*_ record the number of cells within the sample of *n*_samp_ cells that have receptor *σ*. Conditional upon the clonotype sizes, we treat this empirical frequency as Poisson distributed, considering typical experimental settings and the relative rarity of individual clonotypes (e.g., Sepúlveda, Paulino and Carneiro, 2010). Thus,

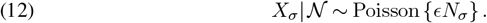

The number of clonotypes represented by *k* cells in the sample is *Y*_*k*_ = ∑_*σ*_ 1[*X*_*σ*_ = *k*]; most diversity statistics are computed from these occupancy counts, {*Y*_*k*_} (e .g., Lande, 1996; Zhang and Zhou, 2010; Chiffelle et al., 2020). The most simple one is 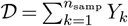, which is the number of distinct clonotypes observed in the sample. Note also *n*_samp_ = ∑_*k*_ *kY*_*k*_ . Recognizing D = ∑_*σ*_ 1[*X*_*σ*_ *>* 0], it is immediate from exchangeability that:

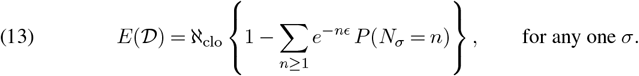

Using characteristic functions, we may compute expected diversity directly for the reference marginals. For example, taking the limiting Geometric margin for *P* (*N*_*σ*_ = *n*) noted in Section 3.2,

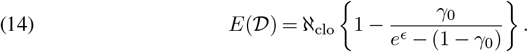

If *N*_*σ*_ ∼ Log(*p*), then,

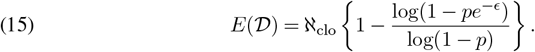

For Yule-Simon marginal distribution with parameter *ρ*, we get,

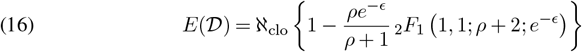

where _2_*F*_1_(*a, b*; *c*; *z*) is the Gaussian hypergeometric function. In typical repertoires, we expect parameter settings assuring high diversity, such that *E*(𝒟) is relatively close to *n*_samp_.

Surrogate selection enables direct sampling from the mutant fraction, and our formalism allows a quantitative assessment of the selection effect on expected sample properties. By enriching for larger clonotypes, surrogate selection would seem to lead to fewer cells from very small clonotypes, and thus less diverse samples. Here we confirm that property. Set 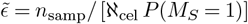, which is an amount larger than *E* that is sufficient to produce, in expectation, *n*_samp_ mutant cells from the repertoire. These cells arise from the clonotypes according to sample counts 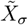, which, given the total numbers of mutant counts across the repertoire, 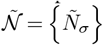, then satisfy

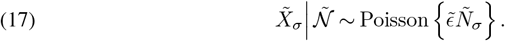

The mutant sample, which in expectation has the same number of mutant cells as the total number of cells in the full-repertoire sample, has its own diversity, 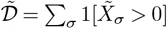. By manipulating the probability generating function of the Luria-Delbrück distribution, and also leveraging results in Roshan, Jones and Greenman (2014), we find explicit formulas for the expected diversity among mutant-sampled cells.

proposition 4. *In the pure-birth, Yule tree model for clonotype development, with a Geometric*(*γ*_0_) *distribution for each clonotype size at observation time, and with mutation frequency θ as in Proposition 1*, *the mutant sample has expected diversity:*

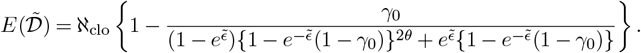

*Alternatively, in case the clonotype-size distribution is* Logarithmic(*p*), *then the expected diversity is:*

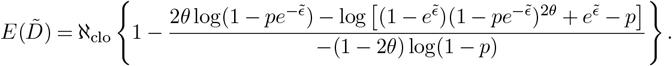

*In either case*, 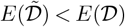 *as long as θ* ∈ (0, *ϵ/*2).

Thus in two reference models, Proposition 4 expresses the precise effect of surrogate selection on repertoire sample diversity; Figure 6 provides a numerical illustration. The result extends to more general distributions by mixing. For example, if conditional upon *γ*_0_ the clonotype sizes are Geometric(*γ*_0_), and if *γ*_0_ = exp(−*W*) for *W* ∼ Exp(*ρ*), then marginally the clonotype size is Yule-Simon distributed with parameter *ρ*, and the expected diversity bound carries through the expectation: 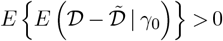.

**FIG 6.**
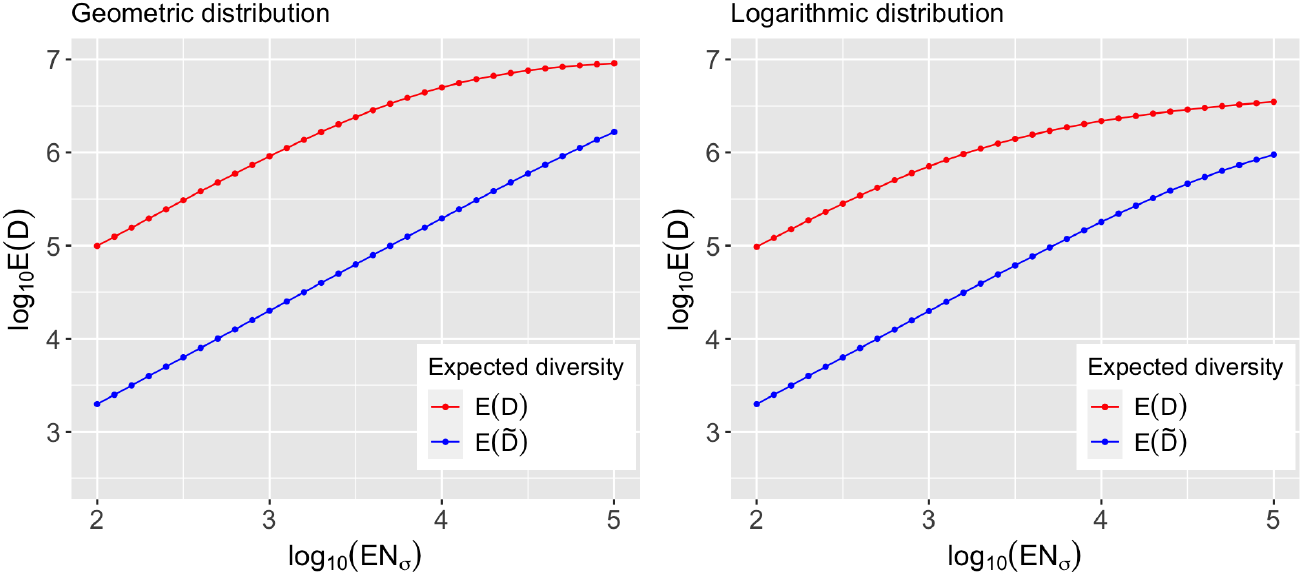
Comparison of expected diversity scores between sampling from whole repertoire or just the mutant fraction, under various Geometric (left) and Logarithmic (right) distributions. The range of Geometric parameter γ_0_ and logarithmic parameter p is determined to match a clonotype of approximately 10^2^ to 10^5^ cells, in expectation. Other parameters are fixed as sampling fraction ϵ = 10^−4^, overall number of clonotypes ℵ_clo_ = 10^7^ and mutation probability in each division θ = 10^−6^. Expected diversity is always lower in the mutant fraction, in line with Proposition 4

### 3.6. Somatic burden

Our calculations emphasize mutation status at some special locus (like HPRT) for which experimental assays provide for ready sampling of cells within that mutant fraction of the repertoire. Yet the calculations also inform an analysis of more general mutational signatures carried by sampled T cells. Intuitively, there may be a lot of information, for example about prior antigen exposure, that is recorded in present genomic state of sampled T cells, whether or not we consider mutations for an *in vitro* selection assay.

A T cell sampled randomly from the repertoire resides in a random clonotype *S* of size *N*_*S*_. At any genomic locus *g* within a host of measurable sites 𝒢, this cell has mutation status *M*_*S*,*g*_ relative to its prethymic state. We are thinking

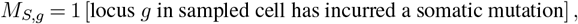

which opens us up to a genome-wide spectrum of mutations, rather than changes at a single, surrogate-selection-driving locus. To this end, we define a sampled cell’s *somatic burden L* to be the summation of *M*_*S*,*g*_ over all *g* ∈ G. We find it convenient to consider a sequence of collections 𝒢^1^, *𝒢*^2^, · · ·), approaching 𝒢, with 𝒢^*m*^ containing *m* loci, and for which at step *m*, 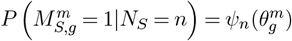 for locus-specific mutation frequency 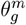, and with *ψ*_*n*_ asin (3) but now highlighting its dependence on mutation frequency. This formula works in the pure-birth model structure thanks to Proposition 1 and the exchangeability in (8). Within this framework, we have the step-*m* burden 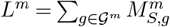.

proposition 5. *If clonotypes satisfy the regularity conditions in Proposition 1*, *if clonotype sizes are exchangeable as in (8)*, *and if* 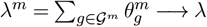 *as m* → ∞ *for some λ >* 0, *then*

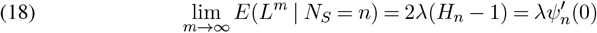

*where H*_*n*_ *is the n*^*th*^ *harmonic number and* 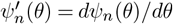.

Put another way, the expected number of post-thymic somatic mutations in a T cell is approximately proportional to the logarithm of that cell’s clonotype size, at least under the stated regularity conditions. Single-cell sequencing studies provide a means to measure *L* on sampled cells, and also to associate that somatic burden with clonotype size, as we investigate next.

## 4. Empirical studies

### 4.1. Somatic burden

Single-cell sequencing technologies provide an exciting window into the dynamics of the T cell repertoire. Here we reanalyze publicly available data reported by 10x Genomics on samples from 7 different T cell repertoires, including 5 peripheral blood mononuclear cell (PMBC) samples from healthy human donors, a melanoma patient and a lung cancer patient. Supplementary Material, Appendix F, summarizes the data resources and provides additional details on our analysis pipeline. In every case, the repertoire sampling and prior analysis provided both the T cell receptor (TCR) sequence and single cell whole-transcriptome RNA-seq on thousands of cells. The TCR sequence information allows us to cluster cells into clonotypes. Our interest in somatic burden puts quite different demands on the RNA-seq data than the original studies. Rather than derive transcript abundance, we repurpose the RNA-seq reads to report on underlying somatic mutations that must have emerged in the genomic DNA. Following the workflow in Edwards et al. (2022), and using the GATK pipeline for genomic-variant calling (McKenna et al., 2010; Auwera and O’Connor, 2020), we computed single-cell-expressed single-nucleotide-variant calls (sce-SNVs) from the aligned read data using Mutect2 (Cibulskis et al., 2013; DePristo et al., 2011), applied consistently across the different repertoires. Details for SNV calling are in Appendix F, but we note here that to focus better on post-thymic somatic variants, we filtered any calls that would have appeared in more than one clonotype. In total over the 7 repertoires, we measured 30257 cells that resided in 27758 clonotypes, and which altogether presented 1609 post-thymic sce-SNVs.

Figure 7 summarizes average somatic burden as a function of clonotype size for one repertoire. Though not statistically significant, it shows an intriguing increase in estimated mean burden with increasing clonotype size, just as predicted by Proposition 5. Not all data sets show as clear a trend (Table 1), though in a meta-analysis which combines the 7 repertoires, we see stronger evidence of an increase in expected burden with clonotype size. We applied a linear model to cell-level data, with response the measured burden, and with an adjusted clonotype size predictor, where the adjustment accounts for the different sampling rates across the repertoires. We estimate 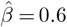 SNVs per unit increase in logarithm of clonotype size. A stratified permutation, which shuffles cells between clonotypes within repertoires, gives a modest p-value of 0.02 on this clonotype-size effect. Further details are in Figure S5.

**Table 1.** Somatic burden of cells by clonotype size (rows), derived from seven T cell repertoire samples (columns) made publicly available by 10x Genomics. *Details of the data resources are in Supplementary Table S3*. *We repurposed the single-cell RNA-seq reads to infer somatic variants and compute somatic burden counts per cell (average burden in upper table, SNVs/cell); and we used the reported TCR sequences to partition cells into clonotypes (numbers of clonotypes in bottom table)*.

| Clonotype size | 20K | 10K | SC5K | PBMC3 | Controller | Melanoma | Lung |
| --- | --- | --- | --- | --- | --- | --- | --- |
| 1 | 0.018 | 0.017 | 0.076 | 0.042 | 0.019 | 0.057 | 0.390 |
| 2 | 0.002 | 0.005 | 0.103 | 0.043 | 0.029 | 0.121 | 0.245 |
| 3 | 0 | 0 | 0 | 0 | 0 | 0.035 | 0.407 |
| 4 | 0 | 0 | - | 0.042 | 0 | 0.278 | 0.667 |
| 5 | 0 | 0 | 0 | 0 | 0 | 0 | 0.400 |
| 6 | 0 | 0 | 0 | 2.167 | 0 | - | 0.292 |
| 7 | 0 | - | 0 | 0 | 0 | - | 0.429 |
| 8 | 0 | 0 | 3.000 | 0 | - | - | 0.875 |
| 9 | 0 | - | 0 | - | - | 0 | 0.444 |
| 10 | 0 | - | - | 0 | 0 | - | 0.200 |
| 11 | 0 | - | 0 | - | 0 | 0 | 0.455 |
| 12 | - | 0 | - | - | 0 | 0 | 0.292 |
| 13 | - | - | - | - | - | 0 | - |
| 14 | 0 | - | 0.429 | - | - | 0 | - |
| 17 | - | - | - | 0 | - | - | 0.588 |
| 19 | 0 | - | - | - | - | 0 | - |
| [20, 40] | 0.100 | 0 | - | - | 0 | - | 1.283 |
| > 40 | 0 | - | 0.170 | 0.171 | - | - | - |
| Clonotype size | 20K | 10K | SC5K | PBMC3 | Controller | Melanoma | Lung |
| 1 | 8395 | 4211 | 1643 | 5659 | 4118 | 1097 | 1315 |
| 2 | 239 | 111 | 39 | 278 | 123 | 66 | 108 |
| 3 | 39 | 35 | 8 | 33 | 23 | 19 | 27 |
| 4 | 13 | 6 | - | 6 | 6 | 9 | 12 |
| 5 | 15 | 5 | 1 | 4 | 5 | 3 | 3 |
| 6 | 7 | 2 | 2 | 1 | 1 | - | 4 |
| 7 | 5 | - | 2 | 2 | 2 | - | 2 |
| 8 | 6 | 1 | 1 | 4 | - | - | 1 |
| 9 | 2 | - | 1 | - | - | 1 | 2 |
| 10 | 1 | - | - | 2 | 1 | - | 2 |
| 11 | 2 | - | 1 | - | 1 | 1 | 1 |
| 12 | - | 1 | - | - | 1 | 1 | 2 |
| 13 | - | - | - | - | - | 1 | - |
| 14 | 1 | - | 1 | - | - | 1 | - |
| 17 | - | - | - | 2 | - | - | 1 |
| 19 | 1 | - | - | - | - | 2 | - |
| [20, 40] | 1 | 1 | - | - | 1 | - | 2 |
| > 40 | 1 | - | 1 | 1 | - | - | - |

**FIG 7.**
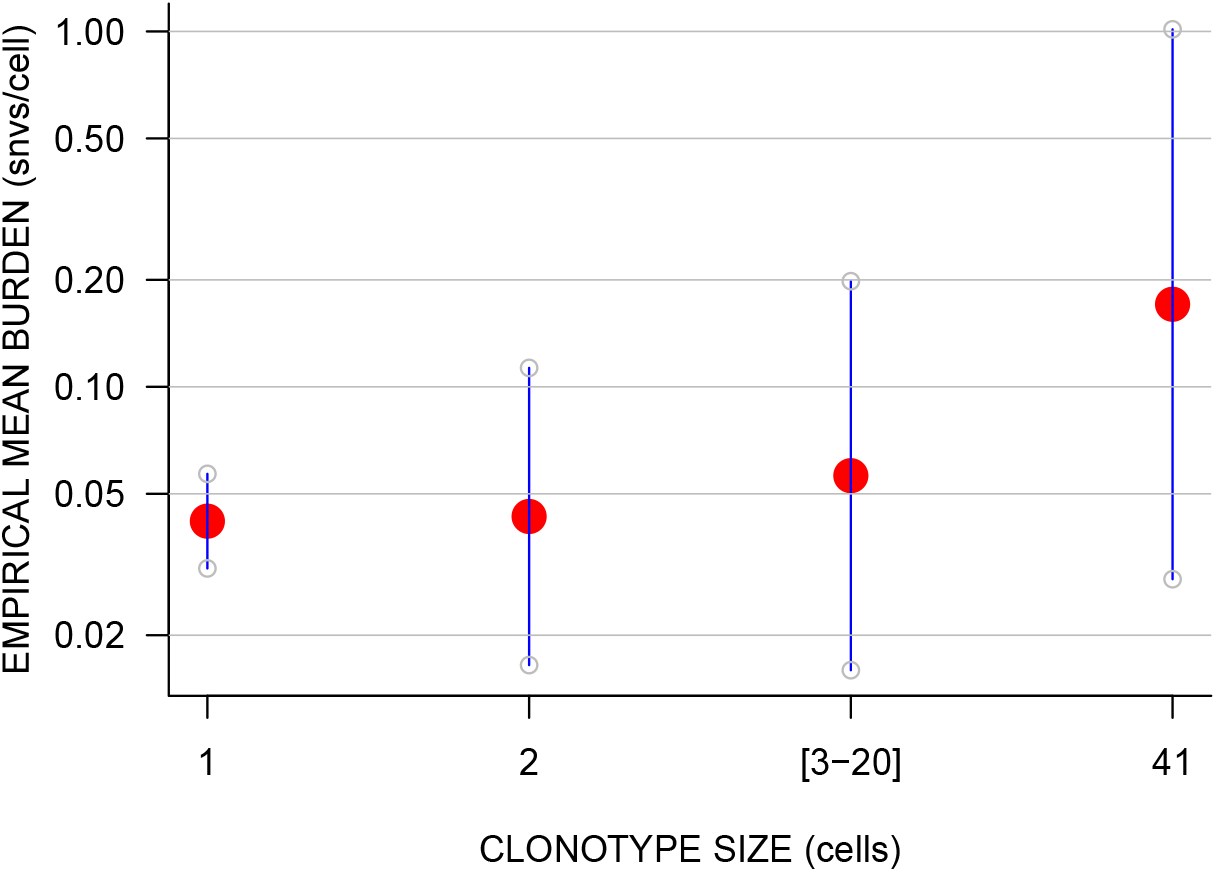
Association of average somatic burden with clonotype size, in the PBMC3 repertoire. There are 5659 singleton clonotypes, 278 duplexes, and a total of 22 clonotypes with sizes greater than 2. The largest clonotype contains 41 cells. Clonotypes of size 3 to 20 cells are combined together as a single class considering the small sample size. Pointwise 95% confidence intervals are computed from a quasi-Poisson generalized linear model.

### 4.2. Melanoma case studies

We reconsider surrogate selection data presented in Zuleger et al. (2020), and we focus here (Table 2) on a metastatic melanoma patient for whom repertoire sampling was performed repeatedly over the course of what turned out to be a successful immunotherapy treatment. As the table shows, the HPRT wild-type (WT) samples have greater sample diversity than the HPRT mutant (MT) samples, which have passed *in vitro* selection.

**Table 2.** Empirical repertoire diversity in wild-type and HPRT mutant fractions, derived from sequencing TCR cDNAs from mass cultures obtained at 5 time-points on one melanoma patient.

| Time point | Total reads | WT unique / reads | MT unique / reads |
| --- | --- | --- | --- |
| 1 | 108722 | 2840 / 58896 | 158 / 49826 |
| 2 | 111652 | 4587 / 53435 | 182 / 58217 |
| 3 | 98834 | 2709 / 49799 | 156 / 49035 |
| 4 | 87804 | 2091 / 52277 | 84 / 35527 |
| 5 | 98286 | 2209 / 51711 | 133 / 46575 |

The mass culture conditions and cDNA sequencing approach used by Zuleger et al. (2020) affect the distribution of counts in Table 2, making them over-dispersed compared to ideal cell counts. Assays based upon single-cell-derived isolates precisely count wild-type and HPRT mutant cells, rather than cDNAs, and are not subject to additional variance caused by in-vitro growth effects. However they are more labor intensive than mass cultures and provide less overall sequencing data. Table 3 summarizes such data from the peripheral blood of 11 subjects studied in Zuleger et al. (2011). In all cases the HPRT surrogate selected samples are less diverse than the wild-type cells, as predicted by the enrichment calculations in Section 3.5.

**Table 3.** Empirical repertoire diversity in wild-type and HPRT mutant fractions, derived from single-cell isolate data on seven melanoma patients and four healthy donors. Subjects 1, 2, 3, 5, 6, 9, 13 are melanoma patiens; Subjects 26, 29, 30, 32 are healthy donors. Subjects are sorted by the number sequenced T cell receptors.

| Subject | # T cells | WT unique / cells | MT unique / cells |
| --- | --- | --- | --- |
| 5 | 122 | 19 / 19 | 102 / 103 |
| 2 | 114 | 49 / 49 | 61 / 65 |
| 1 | 101 | 31 / 32 | 45 / 69 |
| 32 | 95 | 54 / 54 | 30 / 41 |
| 26 | 81 | 36 / 36 | 44 / 45 |
| 3 | 79 | 17 / 17 | 55 / 62 |
| 30 | 69 | 39 / 39 | 29 / 30 |
| 13 | 69 | 23 / 23 | 43 / 46 |
| 29 | 56 | 36 / 36 | 19 / 20 |
| 9 | 50 | 11 / 11 | 23 / 39 |
| 6 | 26 | 18 / 18 | 8 / 8 |

## 5. Concluding Remarks

Gaining a better understanding of the adaptive immune system is a central focus of contemporary biomedical research, considering that system’s role in health and disease. We seek clinically useful methods to identify T cells that may be responding to antigens presented by melanoma, but it is challenging to recognize a patient’s disease-specific antigens, and it is also difficult predict the antigens to which a given T cell receptor will bind. Research on both these frontiers is important and will capitalize on advances in the data sciences (e.g., Lu et al., 2021; Li et al., 2021). In any case, techniques that could readily enrich a lymphocyte sample for T cells responsive to disease-relevant antigens would have a variety of practical applications. The present work provides a statistical basis to the use of surrogate selection, which aims to enrich lymphocyte samples for disease-relevant cells by recognizing that prior clonal expansions may be associated with the accumulation of neutral somatic alterations. Relatively straightforward assays, like HPRT and PIG-A, are available to filter cells having incurred some convenient somatic alteration. Earlier studies have compared selected and unselected cell populations, using both standard and novel statistical tools to account for sources of variation affecting cell phenotypes (e.g., Pei et al., 2014; Zuleger et al., 2020). No prior studies have considered the stochastic basis of surrogate selection itself, and this problem has been the central focus of the present paper.

We treat the stochastic development of a single clonotype and demonstrate that conditioning on a mutant sampled cell enriches for larger clonotypes in a class of birth-death processes (Propositions 1 and 2). We extend the development to exchangeable collections of clonotypes (Proposition 3), accounting for the size bias and complexity of real repertoires. We study the effects of selection on the sampling distribution of a commonly computed diversity statistic (Proposition 4). Looking beyond selection, we investigate the accumulation of neutral somatic mutations across the genome, and show how the same modeling calculations demonstrate that cells in older, expanded clonotypes are expected to carry a greater mutation burden. All these theoretical predictions are accompanied by empirical results both from surrogate selection studies and recent single-cell sequencing projects. If there would be a single take-home message it would be that we have resolved the sampling phenomenon exemplified in the simulated data of Figure 5. Interestingly, cells sampled from this synthetic repertoire are associated with larger clonotypes when we condition on them being mutant, even though mutation events are completely neutral. Moreover, we hope that the quantitative characterizations developed here will provide a basis for more informed statistical analysis of T cell data sets and the planning of immunological experiments.

## SUPPLEMENTARY MATERIAL

We provide derivations, proofs, and additional modeling elements in support of findings presented in the main manuscript, “Surrogate selection oversamples expanded T cell clonotypes”, by Yu, Lian, Zuleger, Albertini, Albertini, and Newton. We also provide further details regarding data preparation and analysis from Section 4 of that work. Supplementary material is organized in seven appendices indicated below; the section in parentheses refers to the numbering within the main manuscript.

## APPENDICES

A: Enrichment in single clonotype case (Sections 2.4 and 2.5)

B: Poisson induced assemblages (Section 3.2)

C: Mutant Frequency (Section 3.4) D: Expected Diversity (Section 3.5)

E: Burden Statistics (Section 3.6) F: Variant calling (Section 4.1)

G: Additional Figures and Tables

## APPENDIX A

ENRICHMENT IN SINGLE CLONOTYPE CASE (SECTIONS 2.4; 2.5)

### Pure birth case

Proposition 1 is established by the arguments in and leading up to Section 2.4. Recall the mutant frequency among cells in a pure-birth clonotype of a given size:

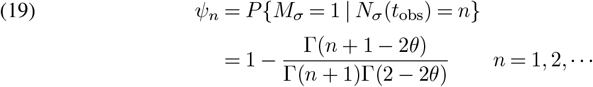

as in Eq (4).

### Birth-death cases

#### Counter example

There are birth-death processes for which surrogate selection fails to enrich the sample for larger clones. Suppose a clonotype *σ* develops by a linear pure birth process up to some time *t*_0_, and consider some fixed threshold *K* ≥ 3. Suppose also that for *t > t*_0_, *N*_*σ*_(*t*) = *N*_*σ*_(*t*_0_) with probability one if *N*_*σ*_(*t*_0_) *< K*, and *N*_*σ*_(*t*) = 1 with probability one if *N*_*σ*_(*t*_0_) ≥ *K*. In other words, the birth rate drops to 0 and the death rate remains 0 after *t*_0_ if the clone size is less than *K*; otherwise the death rate becomes ∞ until exactly one cell survives. This is a highly stylized model in which a clonotype of size less than *K* at time *t*_0_ is essentially naïve; and then after *t*_0_ it is more like a post-activation T cell clone where only one memory T cell remains. Here, the mutant frequency may not increase along with the clone size. For example, at *t*_obs_ *> t*_0_,

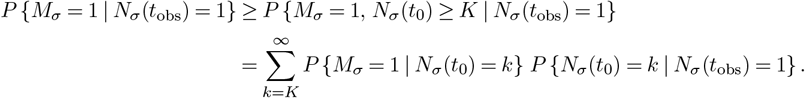

The first factor in each summand does not involve *N*_*σ*_(*t*_obs_) explicitly, since in this event the sampled cell yielding *M*_*σ*_ = 1 is the same as the single surviving cell after the spate of cell deaths. And this factor is exactly the conditional mutant frequency *ψ*_*k*_ for a pure-birth process, which we recall is strictly increasing in its argument. Furthermore, by the structure of the process after *t*_0_, and for *k* ≥ *K*,

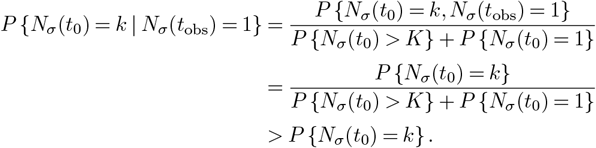

Taking *K* = 3 as a simple case and combining the results above, we get,

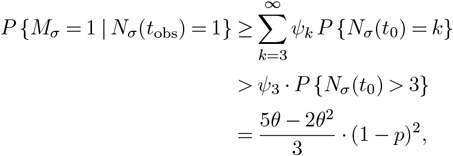

which at *θ* = 0.1 and for Geometric parameter *p* = 0.1 gives the bound 0.1296, for example. On the other hand, given *N*_*σ*_(*t*_obs_) = 2, the mutant frequency is:

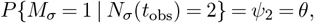

which equals to 0.1 under that parameter setting. Therefore, *P* {*M*_*σ*_ = 1 | *N*_*σ*_(*t*_obs_) = 1} *> P* {*M*_*σ*_ = 1 | *N*_*σ*_(*t*_obs_) = 2} in this toy example. Furthermore, the enrichment ratio *ϕ*_*n*_ could be less than 1 for any feasible clone sizes, as the largest possible clone size is *K* when observation time *t*_obs_ *> t*_0_, and all *ϕ*_*n*_ *<* 1 as long as the threshold is less than the crossover point, i.e. *K < n*_cross_.

proof of proposition 2. For the mutation frequency

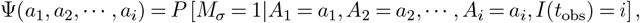

we first establish the useful recursion:

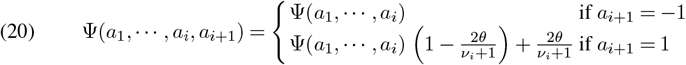

where 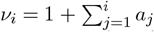 is the clonotype size after *i* steps. To prove (20), it is helpful to introduce *X*_*i*_ recording the number of mutant cells among the *ν*_*i*_ cells in the clonotype just after step *i*: so, 0 ≤ *X*_*i*_ ≤ *ν*_*i*_ − 1, recalling that the originating cell is non-mutant in our model, and new mutants appear as at most one of the two daughter cells. Owing to the sampling of a cell at random to determine *M*_*σ*_,

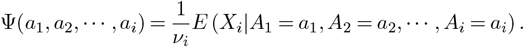

Moving to step *i* + 1, the distribution of *X*_*i*+1_ given past steps and *A*_*i*+1_ depends on the whether *A*_*i*+1_ is a birth (*a*_*i*+1_ = 1) or a death (*a*_*i*+1_ = − 1). In the case *a*_*i*+1_ = 1, *ν*_*i*+1_ = *ν*_*i*_ + 1, and *X*_*i*+1_ = *X*_*i*_ + *C*_*i*+1_ where *C*_*i*+1_ is a Bernoulli trial taking value 1 if the dividing cell is mutant or if the dividing cell is non-mutant but a new mutation emerges from the division. Thus, conditional on *X*_*i*_ and the past sequence of birth-death steps,

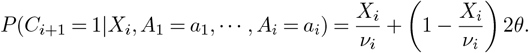

Taking conditional expectations to average over *X*_*i*_,

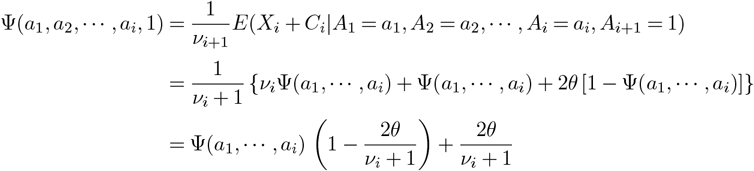

For the death event *a*_*i*+1_ = −1, *ν*_*i*+1_ = *ν*_*i*_ − 1; also *X*_*i*+1_ is *X*_*i*_ if the death is of a non-mutant cell and equals *X*_*i*_ − 1 if the death is of a mutant cell. Taking expectations confirms that Ψ(*a*_1_, …, *a*_*i*_, − 1) = Ψ(*a*_1_, …, *a*_*i*_), and so (20) is established. Intuitively, the removal of a random cell does not change features of the remaining cells.

By convexity of combinations in recursion (20), the random sequence *Z*_*i*_ := Ψ(*A*_1_, *A*_2_, …, *A*_*i*_) is almost surely non-decreasing in *i*. In fact, *Z*_*i*+1_ = *Z*_*i*_ + 1[*A*_*i*+1_ = 1](1 − *Z*_*i*_)(2*θ*)*/*(*ν*_*i*_ + 1), and so

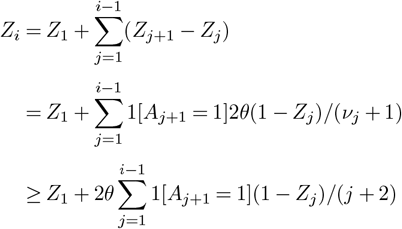

because the clone size *ν*_*j*_ ≤ *j* + 1. Monotonicity also implies that *E*(*Z*_*i*+1_ | *Z*_*i*_) ≥ *Z*_*i*_, which means the sequence forms a submartingale. By the martingale convergence theorem (e.g., Billingsley, 1995, pg 468), *Z*_*i*_ converges almost surely to some limit *Z* ∈ [0, 1], approaching the limit from below, and so for all *j*, 1 − *Z*_*j*_ ≥ 1 − *Z*. Thus

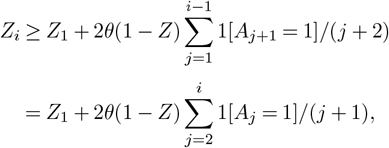

where the second line just invokes a change of variables in the sum. If the limit *Z* is less than 1 on some realization, then 1 − *Z* does not eliminate the subsequent sum. But we had assumed divergence of this sum with probability one, which would create an impossible lower bound for *Z*_*i*_. The only option is for *Z* = 1 with probability one. In other words, the mutation frequency converges to 1 in the general birth-death process as long as there are sufficiently many births.

Convergence of *E*(*Z*_*i*_) is the monotone convergence theorem (e.g., Billingsley, 1995, pg 208).

On the conditional mutant frequency, and with *I* = *I*(*t*_obs_) for shorthand,

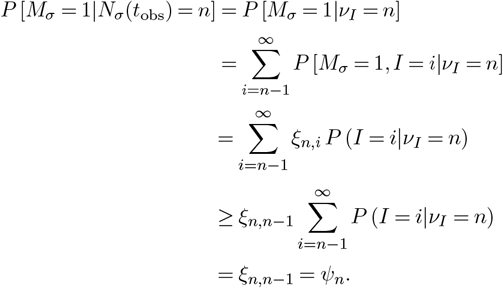

The inequality comes from the assumption that *ξ*_*n*,*i*_ is non-decreasing in *i* for each *n* (e.g. see Fig S1).

We note for clarity that the increasingness of *ξ*_*n*,*i*_ in *i* for each *n* must refer to the values *i* that are allowable in the birth-death formalism. For example, if *n* = 3, then *i* must be in 2, 4, 6, …; in general *i* can range in {*n* − 1, *n* + 1, *n* + 3, …}.

**FIG S1.**
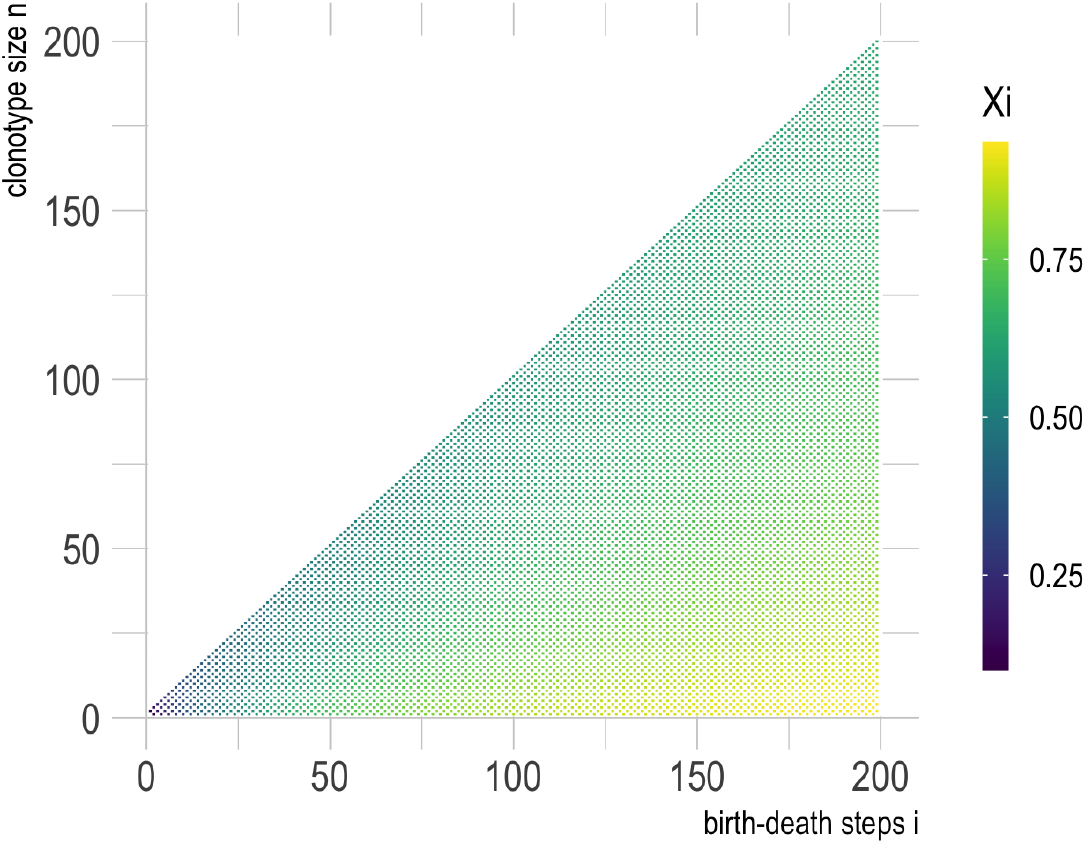
ξ_n,i_ for a linear birth-death process with λ = 19, μ = 1, and θ = 0.01. Numerically, the probabilities increase from left to right over i ∈ {n − 1, n + 1, n + 3, …} for each value n, as required by Prop 2.

## APPENDIX B

POISSON-INDUCED ASSEMBLAGES (SECTION 3.2)

There are several ways in modeling the allocation of ℵ_cel_ T cells among ℵ_clo_ *<* ℵ_cel_ TCR clonotypes. The general assemblage specifications in Rothman and Templeton (1980) are informative and still relatively simple, so we develop them in the present immunological context for completeness. We refer to this class of exchangeable models as Poisson-induced assemblages.

Unconditionally on clonotype sizes or the repertoire size, suppose the whole repertoire is generated from independent Poisson variates, with the Poisson means themselves drawn as i.i.d. from a Gamma mixture distribution. Prior to any conditioning, the resulting clonotype sizes *N*_*k*_, for *k* = 1, 2,, …,ℵ_clo_, are i.i.d. from a Negative Binomial distribution NB(*α, p*), with p.m.f.:

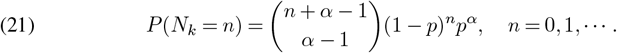

Here, *p* ∈ (0, 1), *α >* 0, and the notation follows Rothman and Templeton (1980), 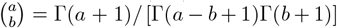. For valid allocations, we condition both on clonotypes being extant and on achieving a certain total repertoire size. That is, with ℵ = (ℵ_clo_, ℵ_cel_) fixed (think of them as very large constants), we will condition on the event

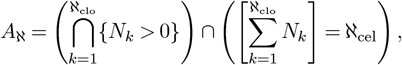

and we consider the induced exchangeable distribution:

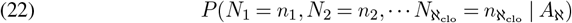

The key to (22) is the probability *P* (*A*_ℵ_), which can be calculated through inclusion-exclusion principle:

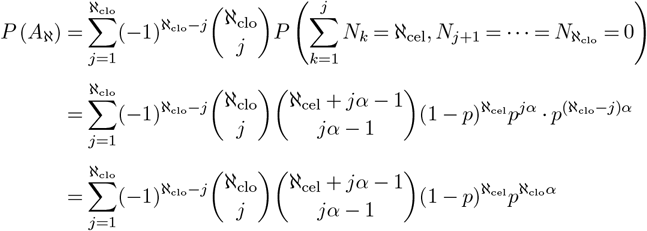

where the second equality is due to the sum of Negative Binomial random variables with same *p* also having a Negative Binomial distribution. Therefore, (22) can be calculated explicitly as:

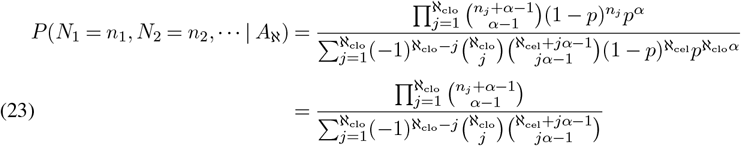

The important special case *α* = 1 is the Bose-Einstein allocation; the distribution in (23) simplifies to

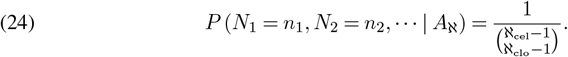

This is because,

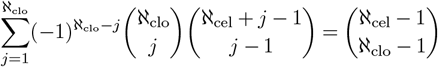

where both sides of the equation count how many ways there are to put ℵ_cel_ indistinguishable balls into ℵ_clo_ distinguishable bins, such that each bin is non-empty. The Bose-Einstein distribution assigns equal probability to all allocations of ℵ_cel_ cells among ℵ_clo_ non-empty clonotypes.

Introducing 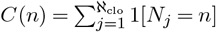 to denote the number of clonotypes comprised of *n* cells, the vector

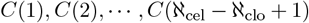

is called the frequency spectrum of the repertoire. These counts-of-counts are sufficient in exchangeable models since (8) depends on the clonotype sizes {*n*_*k*_} only through the frequency spectrum. In the Poisson-induced assemblages considered here,

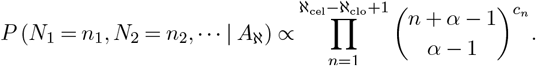

where *c*_*n*_ =∑ _*j*_ 1[*n*_*j*_ = *n*]. By this sufficiency, the probability distribution of the frequency spectrum itself is also proportional to the above. For example, under the Bose-Einstein allocation:

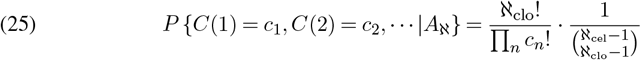

where ∑_*n*_ *c*_*n*_ = ℵ_clo_ and ∑*nc*_*n*_ = ℵ_cel_.

Using these facts about the joint distribution, the size *N*_*σ*_ of any single clonotype follows a Pólya-Eggenberger distribution, PE(1, ℵ_clo_ − 1), if *α* = 1:

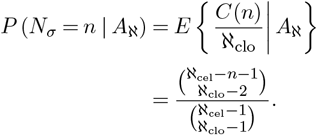

Our interest is on the limiting probability of *P* (*N*_*σ*_ = *n* | *A*_ℵ_) when ℵ_cel_ but ℵ_clo_*/* ℵ_cel_ converges. In the simplest scenario, the limiting ratio is a constant 0 *< γ*_0_ *<* 1, and so by applying Stirling’s formula:

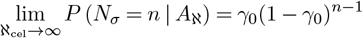

i.e. the distribution of sampled clonotype size converges to a Geometric distribution (on the support *n* = 1, 2,). Noting the original formulation (21) is Geometric when *α* = 1, and on the support *n* = 0, 1, 2, (i.e., including zero), we also see a clear connection between the Bose-Einstein specification (24) and the Geometric margin with non-zero support, and when ℵ_clo_ and ℵ_cel_ are large.

More generally, we may treat ℵ_clo_*/*ℵ_cel_ as random, and suppose it converges in distribution to Θ as ℵ_cel_ diverges. For this case, Hill (1970) proved that under Bose-Einstein allocation,

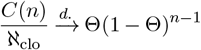

as ℵ_cel_ → ∞. Therefore, the limiting distribution

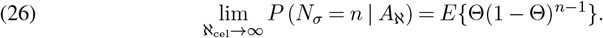

Different cases arise depending on the distribution of Θ in (26). If Θ ∼ Beta(*ρ*, 1), we arrive at the Yule-Simon limit:

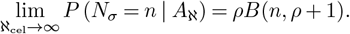

In other words, with Bose-Einstein allocation of ℵ_cel_ cells among ℵ_clo_ clonotypes, if the ratio ℵ_clo_*/* ℵ_cel_ converges to a Beta random variable, then the distribution of sampled clonotype size converges to a Yule-Simon distribution.

The current development allows us to directly calculate the implications of size bias caused by sampling cells from the repertoire. Suppose we sample one T cell uniformly at random from the whole repertoire, and it happens to be from random clonotype *S*. The size *N*_*S*_ of this clonotype satisfies:

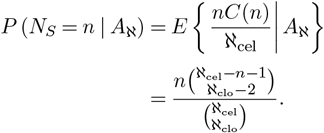

If 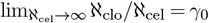 for a positive constant *γ*_0_, then

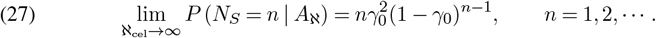

i.e., the limiting distribution of size-biased clonotype size is no longer Geometric, but instead is a Negative Binomial distribution, shifted by 1, that is: 1 + NB(2, *γ*_0_), with support on positive integers. Similarly, if 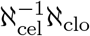 converges in distribution to Θ ∼ Beta(*ρ*, 1) distribution, we find,

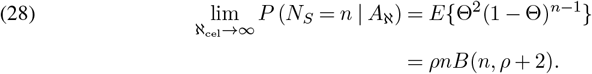

This limit is also a power-law distribution, with

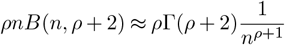

for large *n*. Notice that Yule-Simon distribution p.m.f. *ρB*(*n, ρ* + 1) is approximately *ρ*Γ(*ρ* + 1)*n*^−*ρ*−1^ for large *n*; the size bias effect curiously does not change the tail weight in this case. Figure S2 shows the size-bias effect on Yule-Simon distribution.

Our last comment on Bose-Einstein allocation is its equivalency to Dirichlet-Multinomial distribution DirMult(*n, a*), where 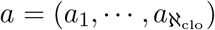 are the (positive) shape parameters of Dirichlet distribution. The DirMult distribution is applied in simulation of Fig 5 (main manuscript). The Bose-Einstein allocation and Dirichlet-Multinomial distribution can both be realized by Pólya-Eggenberger’s urn model: suppose an urn contains ℵ_clo_ balls of distinct colors, at each time a random ball is drawn form the urn and replaced with two balls of the same color. Repeating the sampling and replacement procedure ℵ_cel_ − ℵ_clo_ times will lead to Bose-Einstein allocation with _cel_ balls in _clo_ colors. In the mean time, this Pólya urn model also realizes the DirMult(ℵ_cel_ − ℵ_clo_, 𝟙.) distribution with ℵ_cel_ − ℵ_clo_ trials and all shapes *a*_*i*_ equal to 1. Therefore, the Dirichlet-Multinomial simulation of repertoire in Fig 5 is the same as Bose-Einstein allocation of 1000 cells among 100 clonotypes, which is also the mixture of 100 marginally Geometric-sized clonotypes with 1000 cells in total, as noted previously.

The limiting cases considered above do not handle the case of the Logarithmic marginal. However, we note that taking a limit *α* → 0 in (21), and conditioning on *N*_*k*_ ≥ 1, then the NB distribution converges to the Logarithmic distribution, with p.m.f. proportional to *p*^*n*^*/n*, as in (Kendall and Stuart, 1977, page 139). Further, the joint distribution of the frequency spectrum becomes the Ewens sampling formula (Tavaré, 2021).

**FIG S2.**
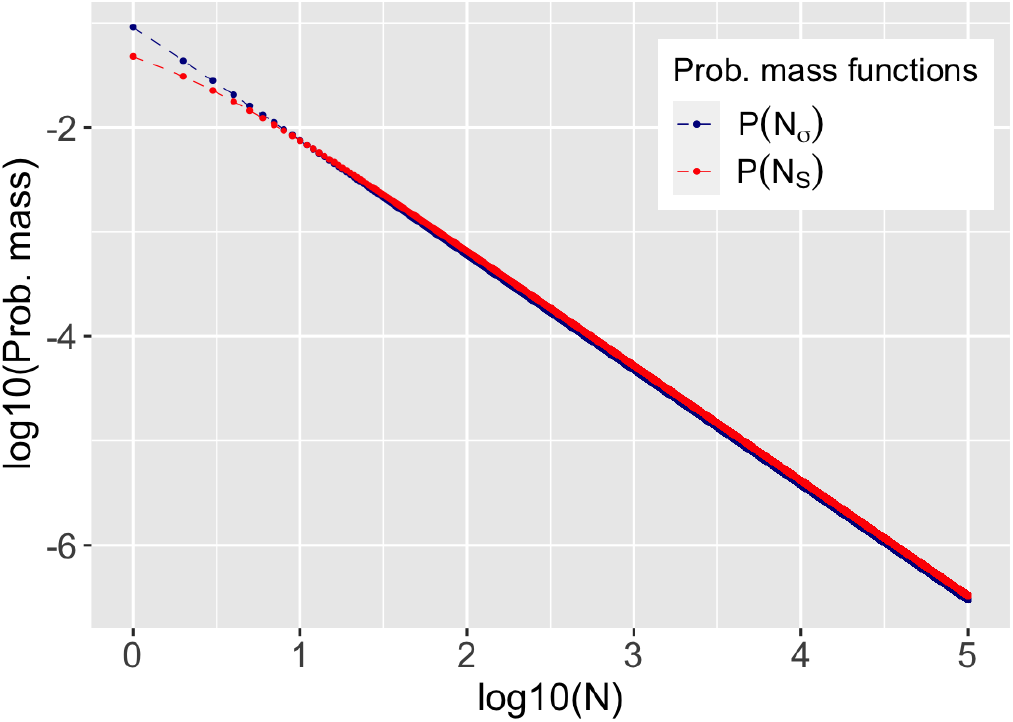
Comparison of limiting distributions of clonotype sizes between size-biased sampling and uniform sampling. Here uniform sampling means uniformly sample a clonotype from the repertoire, while size-biased sampling means uniformly sample a cell from the repertoire.

## APPENDIX C

MUTANT FREQUENCY (SECTION 3.4)

First we show that size bias inflates *P* (*M*_*S*_ = 1) over *P* (*M*_*σ*_ = 1) for any clonotype *σ*. We work with a finitely exchangeable joint assemblage, such as in Appendix B. Recall that *ψ*_*n*_ = *P* (*M*_*σ*_ = 1 | *N*_*σ*_ = *n*) is the mutant frequency defined by Luria-Delbrück distribution, as in (19). Expanding the marginal probability and using exchangeability,

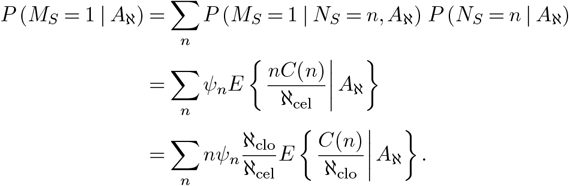

The inflation of mutant frequency is reflected in the term *nψ*_*n*_ in the last equation, as surrogate selection will further enrich the mutants in larger clonotypes. In the Bose-Einstein case, for example,

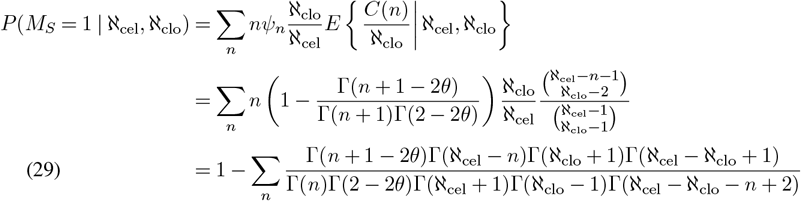

where the second equation is from the Bose-Einstein allocation of ℵ_cel_ cells among ℵ_clo_ clonotypes. The summation in (29) is finite from 1 to ℵ_cel_ − ℵ_clo_ + 1, and hence we can directly calculate the mutant frequency of the repertoire condition on ℵ_cel_ and ℵ_clo_. In reality, ℵ_cel_ ranges from 10^9^ to 10^10^ in the blood, and ℵ_clo_ ranges from 10^6^ to 10^8^. Table S2 compares the marginal mutant frequency for various mutation probability *θ* and number of clonotypes ℵ_clo_, with total number of T cells fixed at ℵ_cel_ = 10^9^. The calculated mutant frequency will decrease as *θ* decreases or average clonotype size ℵ_cel_*/*ℵ_clo_ decreases.

## APPENDIX D

EXPECTED DIVERSITY FOR LOGARITHMIC AND YULE-SIMON (SECTION 3.5)

The expected diversity can be derived directly from the characteristic function of marginal distributions, noting (13). For Logarithmic marginal distribution, *N*_*σ*_ ∼ Log(*p*), we find:

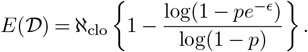

For Yule-Simon marginal distribution with parameter *ρ*, the expected diversity is more complicated. We calculate:

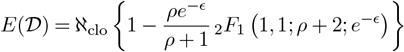

where _2_*F*_1_(*a, b*; *c*; *z*) is the Gaussian hypergeometric function.

proof of proposition 4. From definition of diversity statistic 𝒟,

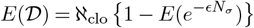

When marginally *P* (*N*_*σ*_ = *n*) = (1 − *γ*_0_)^*n*−1^*γ*_0_, we have

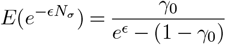

from the characteristic function of the Geometric distribution, and hence

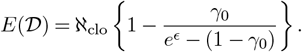

Similarly, the key to 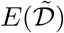 is the expectation 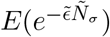. We refer to Theorem 3.2 in Roshan, Jones and Greenman (2014) to derive this expectation by conditioning on *N*_*σ*_. That paper derives an explicit formula for the generating function of (*N*_*σ*_, *W*_*σ*_), where *N*_*σ*_ is the size of the entire clonotype and *W*_*σ*_ is the number of wild type cells. These sizes are defined for extant clonotypes, i.e. *N*_*σ*_ ≥ 1. The generating function used is:

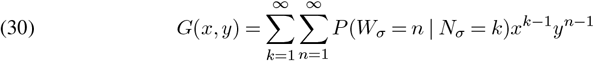

If we let 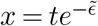 and 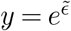 in Eq. (30), we have:

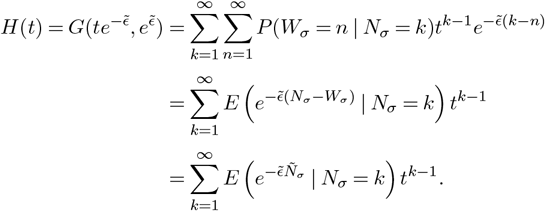

The last equation is because of equality *Ñ*_*σ*_ = *N*_*σ*_ − *W*_*σ*_. If *N*_*σ*_ ∼ Geom(*γ*_0_), we can let *t* = 1 − *γ*_0_ for *H*(*t*) to get the explicit formula for 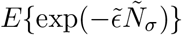:

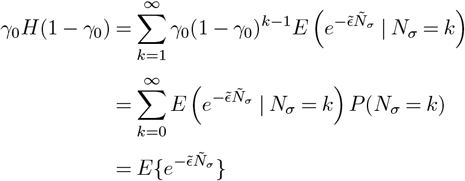

From Theorem 3.2 in Roshan, Jones and Greenman (2014), there is an explicit formula under Luria-Delbrück distribution:

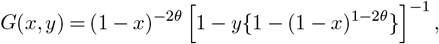

which leads to the formula of *H*(*t*):

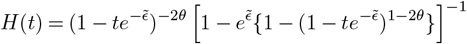

and hence we can get 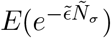:

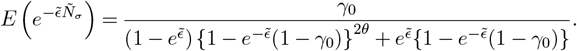

For the second part of Proposition 4, the diversity calculation depends on a comparison of drop-out probabilities. Define

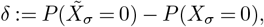

which we see from the definition of our simple diversity statistic satisfies 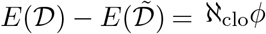, and so it suffices to show that *δ >* 0. From Poisson sampling,

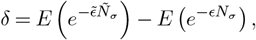

To prove *δ >* 0 for *N*_*σ*_ ∼ Geom(*γ*_0_), we need to show that:

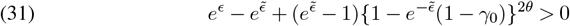

Since *γ*_0_ ∈ (0, 1) and the left-hand-side of (31) is increasing as *γ*_0_ increases, Eq. (31) is equivalent to:

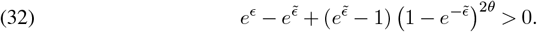

Since function *f* (*x*) = −*x* + (*x* − 1)(1 − 1*/x*)^2*θ*^ is a decreasing function on *x* ∈ (1, *e*) and 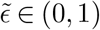, it can be shown that:

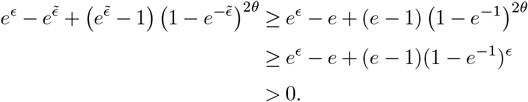

where the second inequality is from the condition *θ < ϵ/*2 and the last inequality is due to the infimum is achieved at *ϵ* = 0. Therefore, we have proved that *δ >* 0 and hence 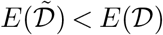 when *N*_*σ*_ ∼ Geom(*γ*_0_).

In another case, if *N*_*σ*_ follows a Logarithmic distribution instead, with p.m.f.

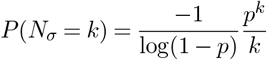

for parameter *p* ∈ (0, 1), then the desired expectation 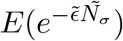 is:

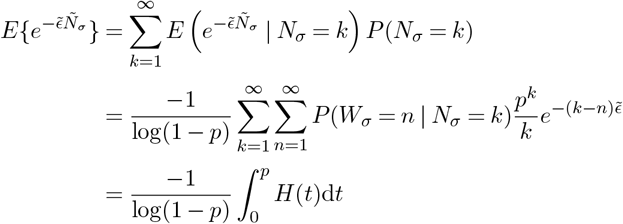

where 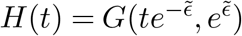 is previously defined. The integral in last equality is valid since the singularity of *H*(*t*) is at:

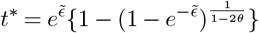

which satisfies *t*^*^ *>* 1 *> p* for 0 *< θ < ϵ/*2 and 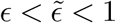. Therefore, the expectation under Logarithmic distribution is:

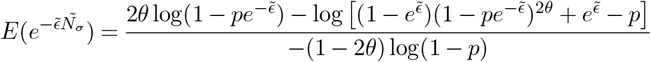

We now show that inequality 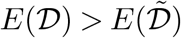 also holds for logarithmic distribution, i.e.,

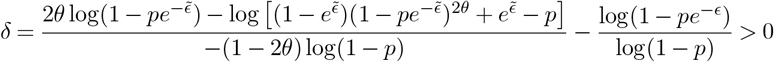

It is equivalent to:

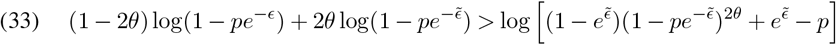

Since 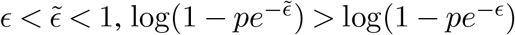, and hence (33) is true if we prove:

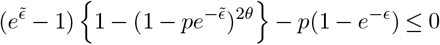

Notice that function *f* (*x*) = (*x* − 1) {1− (1 − *q/x*)^2*θ*^ is an increasing function for 1 *< x < e*, therefore above inequality is equivalent to:

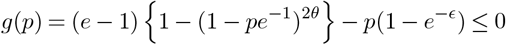

With condition 0 *< θ < ϵ/*2, the second order partial derivative 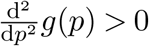 for any 0 *< p <* 1, therefore the supremum of *g*(*p*) is achieved at *p* = 0 or 1. Since *g*(0) = 0 and *g*(1) ≤ 0, it is guaranteed that *g*(*p*) ≤ 0 for 0 *< p <* 1. Therefore, our desired inequality (33) is true, and hence 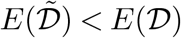.

## APPENDIX E

BURDEN STATISTICS (SECTION 3.6)

proof of proposition 5. With 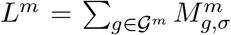 (and within one clonotype *σ*, since we are fixing clonotype size at *n*), we know that 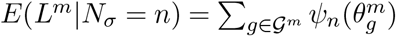, by the findings in Section 2, where 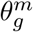 is the mutation frequency associated with locus *g* among the multitude in 𝒢^*m*^. These 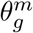’s must be fairly small, considering the convergence of 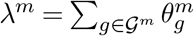. Thus by a Taylor expansion around *θ* = 0,

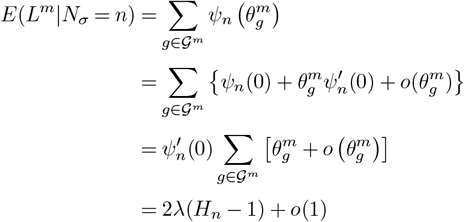

noting 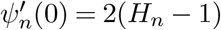 is the derivative of *ψ*_*n*_(*θ*) at *θ* = 0.

## APPENDIX F SOMATIC VARIANTS IN T CELLS FROM SCRNA-SEQ DATA

Our numerical experiments used seven publicly available 10x Genomics data sets obtained from https://www.10xgenomics.com/resources/datasets; the basic characteristics of these data are in Table S3. The 10x Genomics workflow generates scRNA and TCR sequencing reads by capturing and lysing single cells in Gel Beads-in-emulsion which contain a unique oligonucleotide barcode in each bead. According to the reported protocol, mRNA from lysed cells was reverse transcribed to barcoded cDNA. The cDNA was then PCR amplified to construct 5’ Gene Expression library and TCR library. The TCR library consists of V(D)J segments amplified using TCR region specific primers. Sequences were then obtained from the two libraries by Illumina sequencing. Further, the clonotypes of T cells were identified by 10x Genomics by the expressed TRA and TRB genes from TCR-seq. The clonotype information of the T cells from the ‘VDJ TCR -all contig annotations’ CSV files in the online database were imported for our calculations.

For variant calling, we re-processed the single-cell RNA-seq data. The genome-aligned gene expression BAM files downloaded from the 10x Genomics database are used to call somatic variants by Mutect2 with a workflow adapted from Edwards et al. (2022). The BAM files were first processed by the GATK (v.4.2.6.1) module AddOrReplaceReadGroups (McKenna et al., 2010; Auwera and O’Connor, 2020). The module SplitNCigarReads then split the aligned RNA reads which include N elements because of the RNA splicing events. The reference sequences input in GATK modules are the same as versions used for alignment by the Cell Ranger pipeline, available on 10x Genomics and the Ensemble database. After that, somatic variants were called by Mutect2 (Cibulskis et al., 2013; DePristo et al., 2011) using the processed sequencing reads from all cells. We ran Mutect2 under the tumor-only mode. The public Panel of Normal was input to correct for technical artifacts. A population germline resource was also provided for Mutect2 so that the same alleles as in germline resource were not called somatic variants. The variants in the output VCF files were filtered to only keep the confident single nucleotide variants using BCFtools (v.1.15.1) from SAMtools (Danecek et al., 2021) with parameters TYPE=“snp”, INFO/DP (read depth) *>* 20 and MMQ (median mapping quality by allele) *>* 50. The filtered variants called from all RNA-seq data were then assigned to each single cell with the program vartrix (v.1.1.22) by re-aligning single-cell barcoded reads to each variant locus. Two matrices, each of numbers of reference genotype reads and alternative genotype reads corresponding to cell barcodes, are generated by setting the “scoring-method” to “coverage” mode. These data were post-processed in R to associate somatic burden with clonotype sizes.

## APPENDIX G

ADDITIONAL FIGURES AND TABLES

**FIG S3.**
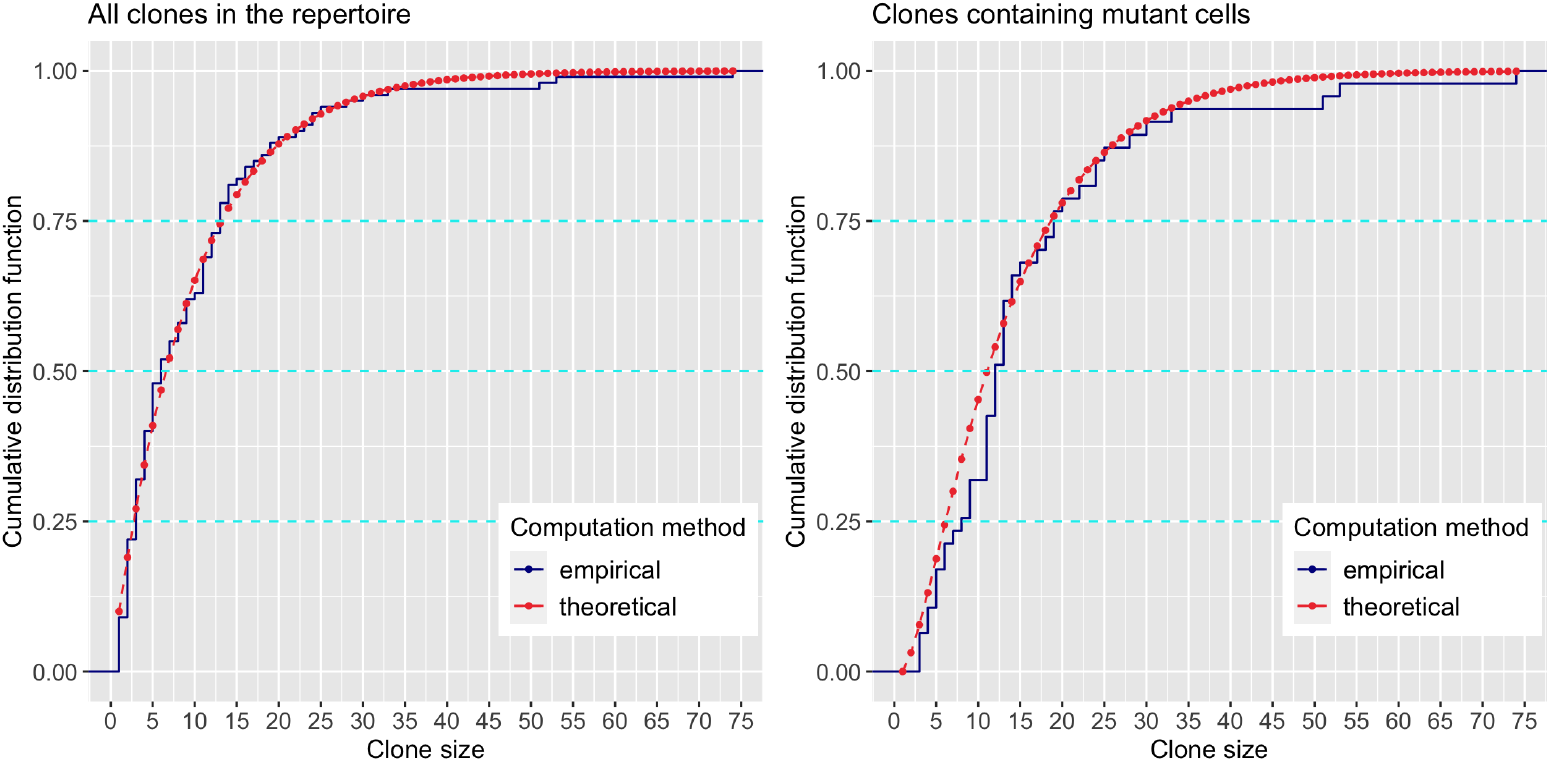
Comparison of empirical and theoretical cumulative distribution functions (c.d.f.) for simulated repertoire in Fig 5. The theoretical c.d.f is calculated from geometric marginal distribution on clonotype sizes, with parameter p = 0.1 and mutant frequency θ = 0.05. These parameters match the repertoire in Fig 5. The solid dark blue lines are empirical c.d.f’s, while dashed red lines are theoretical c.d.f’s. Dashed light blue lines are three quantiles, on which clonotypes with mutant cells are 1.5 to 2 times greater than clonotypes sampled from the whole repertoire.

**Table S1.**
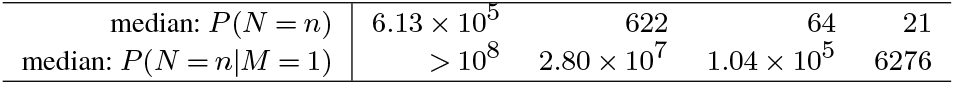
Median clonotype size in the entire repertoire, P (N_σ_ = n), compared to the selected repertoire,P (N_σ_ = n|M_σ_ = 1), for several parametric settings, when θ = 10^−4^

**Table S2.** Mutant frequency P (M_S_ = 1 | ℵ_cel_, ℵ_clo_) in blood samples computed for hypothetical repertoires with various ℵ_clo_ and θ settings. Total cells in the repertoire ℵ_cel_ is fixed at 10^9^.

| $\aleph_{\text{clo}}$ | mutation frequency $\theta$ | | |
| --- | --- | --- | --- |
| | $5 \times 10^{-7}$ | $10^{-6}$ | $5 \times 10^{-6}$ |
| $10^6$ | $7.88 \times 10^{-6}$ | $1.41 \times 10^{-5}$ | $6.93 \times 10^{-5}$ |
| $5 \times 10^6$ | $7.05 \times 10^{-6}$ | $1.33 \times 10^{-5}$ | $5.56 \times 10^{-5}$ |
| $10^7$ | $2.71 \times 10^{-6}$ | $7.42 \times 10^{-6}$ | $4.41 \times 10^{-5}$ |

**Table S3.**
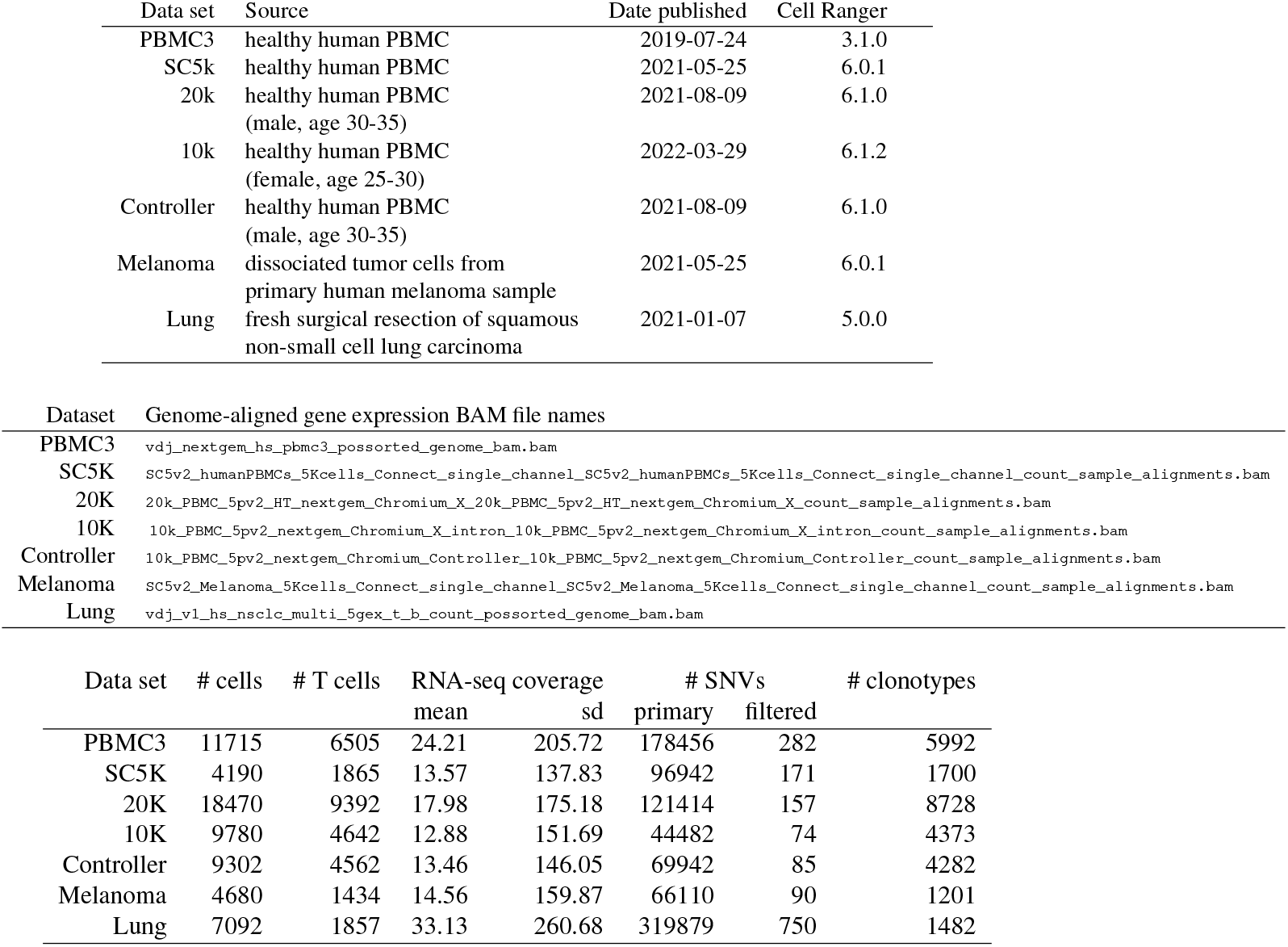
Repertoire sample characteristics. All samples are from human tissue; PBMC=peripheral blood mononuclear cells; data were downloaded from https://www.10xgenomics.com/resources/datasets between June and November, 2022.

**FIG S4.**
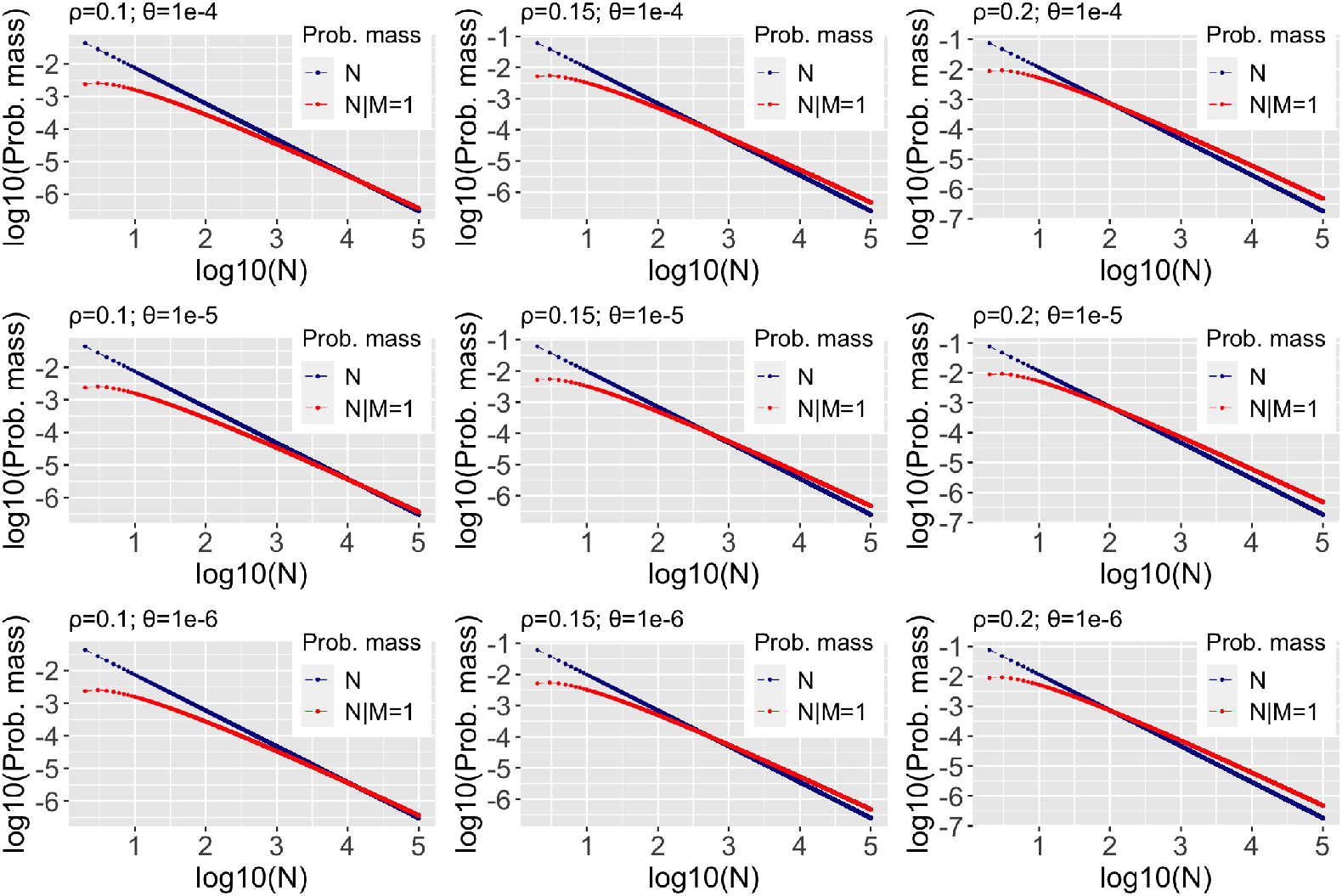
P (N_σ_ = n| M_σ_ = 1) (red) for various power laws P (N_σ_ = n) (blue), on the double log scale.

**FIG S5.**
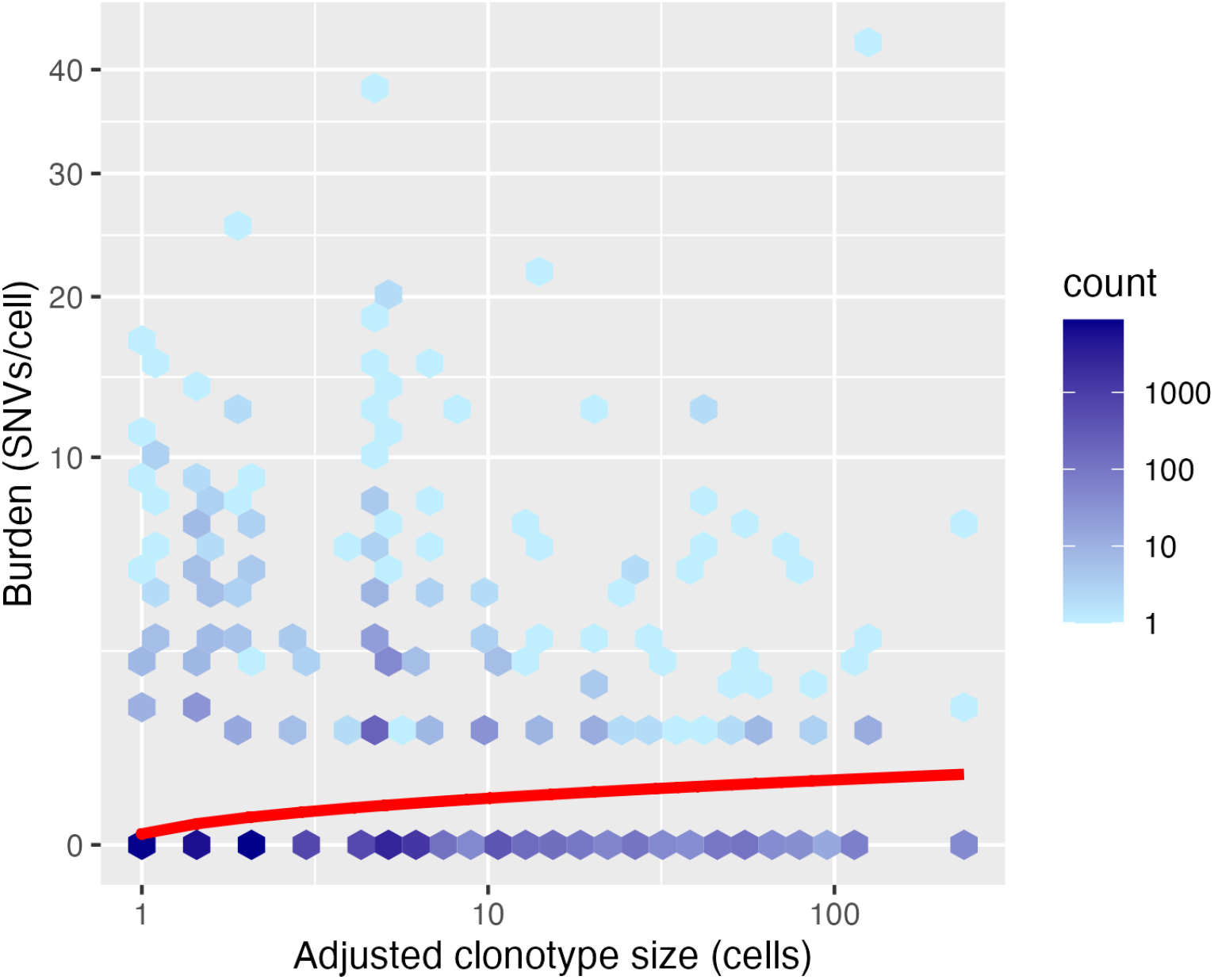
Each of 30257 T cells from 7 repertoires is associated with a somatic burden (vertical) and also a clonotype size (horizontal), the latter of which is adjusted in an effort to normalize repertoire samples. The red curve shows the estimated effect on expected burden of the logarithm of clonotype size, as determined by a linear model fit (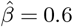 SNVs per unit increase in log clonotype size). Statistical significance of the estimated slope was assessed by a stratified randomization, which shuffled cells between clonotypes but within repertoires (permutation p-value 0.02 with B = 10^4^ shuffles). Though statistically significant, the result is not fully resistant; for example the cells in large clonotypes have very high leverage; dropping the cells with adjusted clonotype size greater than 100, for example, leads to an insignificant permutation p-value. The adjusted log size is log clonotype size minus log of repertoire size plus log of largest repertoire size.

## Footnotes

1 In Mahmoud (1992), a binary tree is assumed to contain *n* internal nodes, and Eq. (2.4) cares about the *n* + 1 external nodes (leaves) of the corresponding extended binary tree. In Steel and McKenzie (2001), following Mahmoud (1992), the Yule tree is said to contain *n* + 1 leaves. Our notation is slightly different as we use *n* to denote leaf numbers. In our setting *n* ≥ 1 and *D*_*σ*_ ≥ 0.

